# Sox2 and FGF20 interact to regulate organ of Corti hair cell and supporting cell development in a spatially-graded manner

**DOI:** 10.1101/436790

**Authors:** Lu M. Yang, Kathryn S.E. Cheah, Sung-Ho Huh, David M. Ornitz

**Affiliations:** Department of Developmental Biology; Washington University School of Medicine; St. Louis, Missouri, 63110; USA; School of Biomedical Sciences; The University of Hong Kong; Pokfulam, Hong Kong, China; Holland Regenerative Medicine Program, and the Department of Neurological Sciences; University of Nebraska Medical Center; Omaha, Nebraska, 68198; USA

## Abstract

The mouse organ of Corti develops in two steps: progenitor specification and differentiation. Fibroblast Growth Factor (FGF) signaling is important in this developmental pathway, as deletion of FGF receptor 1 (*Fgfr1*) or its ligand, *Fgf20*, leads to the loss of hair cells and supporting cells from the organ of Corti. However, whether FGF20-FGFR1 signaling is required during specification or differentiation, and how it interacts with the transcription factor Sox2, also important for hair cell and supporting cell development, has been a topic of debate. Here, we show that while FGF20-FGFR1 signaling functions during progenitor differentiation, FGFR1 has an FGF20-independent, Sox2-dependent role in specification. We also show that a combination of reduction in *Sox2* expression and *Fgf20* deletion recapitulates the *Fgfr1*-deletion phenotype. Furthermore, we uncovered a strong genetic interaction between *Sox2* and *Fgf20*, especially in regulating the development of hair cells and supporting cells towards the basal end and the outer compartment of the organ of Corti. To explain this genetic interaction and its effects on the basal end of the organ of Corti, we provide evidence that decreased *Sox2* expression delays specification, which begins at the organ of Corti apex, while *Fgf20*-deletion results in premature onset of differentiation, which begins near the organ of Corti base. Thereby, *Sox2* and *Fgf20* interact to ensure that specification occurs before differentiation towards the cochlear base. These findings reveal an intricate developmental program regulating organ of Corti development along the basal-apical axis of the cochlea.

**Author summary:** The mammalian cochlea contains the organ of Corti, a specialized sensory epithelium populated by hair cells and supporting cells that detect sound. Hair cells are susceptible to injury by noise, toxins, and other insults. In mammals, hair cells cannot be regenerated after injury, resulting in permanent hearing loss. Understanding genetic pathways that regulate hair cell development in the mammalian organ of Corti will help in developing methods to regenerate hair cells to treat hearing loss. Many genes are essential for hair cell and supporting cell development in the mouse organ of Corti. Among these are *Sox2*, *Fgfr1*, and *Fgf20*. Here, we investigate the relationship between these three genes to further define their roles in development.

Interestingly, we found that *Sox2* and *Fgf20* interact to affect hair cell and supporting cell development in a spatially-graded manner. We found that cells toward the outer compartment and the base of the organ of Corti are more strongly affected by the loss of *Sox2* and *Fgf20*. We provide evidence that this spatially-graded effect can be partially explained by the roles of the two genes in the precise timing of two sequential stages of organ of Corti development, specification and differentation.

## Introduction

The inner ear contains six sensory organs required for the senses of hearing and balance. The cochlea, a snail-like coiled duct, is the auditory organ. It contains specialized sensory epithelia, called the organ of Corti, composed of hair cells (HCs) and supporting cells (SCs). In mammals, this sensory epithelium is elegantly patterned, with one row of inner hair cells (IHCs) and three rows of outer hair cells (OHCs), separated by two rows of pillar cells forming the tunnel of Corti. Each row of OHCs is associated with a row of supporting cells called Deiters’ cells. Here, we refer to pillar cells and Deiters’ cells collectively as SCs.

Organ of Corti development has been described as occurring in two main steps: prosensory specification and differentiation [1]. During prosensory specification, proliferative progenitors at the floor of the developing cochlear duct are specified and exit the cell cycle to form the postmitotic prosensory domain. Here, we define specification to be a process that makes progenitors competent to differentiate. We also use cell cycle exit as a marker for specified cells in the prosensory domain (prosensory cells). During differentiation, prosensory cells differentiate into both HCs and SCs [2]. Interestingly, cell cycle exit, marking the completion of specification, and initiation of differentiation occur in waves that travel in opposite directions along the length of the cochlear duct. At around embryonic day 12.5 (E12.5) in the mouse, progenitors begin to exit the cell cycle and express the cyclin-dependent kinase inhibitor CDKN1B (p27^Kip1^) in a wave that begins at the apex of the cochlea (the cochlear tip) and reaches the base of the cochlea by around E14.5 [3,4]. Afterwards, the specified prosensory cells begin differentiating into HCs and SCs in a wave that begins at the mid-base at around E13.5, and spreads quickly to the rest of the base and to the apex over the next few days [1]. Notably, while the basal end of the cochlear duct differentiates immediately after prosensory specification, the apical end has a longer time between specification and differentiation, providing a larger “temporal buffer” for apical development. The spiral ganglion, containing neurons synapsing to HCs, has been shown to be important for this delay in apical differentiation, via Sonic Hedgehog (SHH) signaling [5–8].

The transcription factor *Sox2* is one of the earliest markers of prosensory cells [9,10]. Mice with specific *Sox2* hypomorphic mutations that affect inner ear expression have hearing impairment due to decreased HC and SC number, while mice with inner ear-specific *Sox2* null mutations are completely deaf and have no HCs or SCs [11,12]. Genetic experiments show that *Sox2* is both necessary and sufficient for prosensory specification. Absence of *Sox2* expression leads to the loss of *Cdkn1b* expression at E14, a marker for the prosensory domain [12], while ectopic *Sox2* expression in cochlear nonsensory epithelium can induce ectopic sensory patches [13–15].

The Fibroblast Growth Factor (FGF) signaling pathway also plays vital roles in organ of Corti development [16]. Studies utilizing cochlear explants showed that inhibition of FGF signaling prior to and during stages of HC and SC differentiation results in decreased HC and SC number [17]. Signaling through FGF receptor 1 (FGFR1), in particular, is essential during this process. Conditional deletion of *Fgfr1* (Fgfr1-CKO) in the developing cochlear epithelium resulted in dramatically reduced HC and SC number [18–20]. This has been attributed to decreased *Sox2* expression in the prosensory domain of Fgfr1-CKO mice, leading to a defect in prosensory specification [19].

FGF20 has been hypothesized to be the FGFR1 ligand during organ of Corti development. Both *in vitro* inhibition of FGF20 with an anti-FGF20 antibody [17] and *in vivo* knockout of *Fgf20* (Fgf20-KO) [21] led to decreased HC and SC number, similar to the Fgfr1-CKO phenotype. However, the Fgf20-KO phenotype is clearly not as severe as that of Fgfr1-CKO. Almost all OHCs and some IHCs are missing in Fgfr1-CKO mice [19], while only 2/3 of OHCs are missing in Fgf20-KO mice, without any loss of IHCs [21]. This suggests that another FGF ligand may be redundant with and compensating for the loss of FGF20, the identity of which is currently unknown.

Another difference between Fgfr1-CKO and Fgf20-KO mice is the proposed mechanism accounting for the decrease in HCs and SCs. Interestingly, unlike in Fgfr1-CKO mice, Sox2 expression in the prosensory domain is not disrupted in Fgf20-KO mice [19,21]. Rather, FGF20 seems to function during HC and SC differentiation. These differences between the Fgfr1-CKO and Fgf20-KO phenotypes and their relationship with *Sox2* suggest that FGF20/FGFR1 signaling has a more complex and as yet unexplained role during organ of Corti development.

Here, we hypothesize that FGFR1 signaling has functions in both steps of organ of Corti development: an earlier role in prosensory specification that involves *Sox2*, and a later role in the initiation of differentiation. We provide evidence that FGF20 regulates differentiation but not specification. Moreover, while *Fgfr1* functions upstream of *Sox2*, *Fgf20* is downstream of *Sox2*. We further show that *Sox2* and *Fgf20* genetically interact during organ of Corti development. Interestingly, downregulation of both genes leads to the loss of hair cells and supporting cells preferentially towards the outer compartment and the basal end of the cochlear duct. To explain the more severe basal phenotype, we provide evidence that *Sox2* regulates the timing of prosensory specification, while *Fgf20* regulates the timing of differentiation. As these two steps occur along a developmental pathway, we hypothesize that prosensory specification must occur prior to differentiation. In *Sox2* hypomorphic mice, prosensory specification is delayed, while in Fgf20-KO mice, onset of differentiation occurs prematurely. When combined, these two defects led to differentiation attempting to initiate prior to the completion of specification towards the basal end of the cochlear duct. These results define unique functions of and complex interactions among FGF20, FGFR1, and Sox2 during organ of Corti development and highlight the potential importance of the timing of specification and differentiation along different regions of the cochlear duct.

## Results

### The Fgf20-KO cochlear phenotype is less severe than the Fgfr1-CKO phenotype

Previous studies showed that deletion of *Fgf20* leads to a loss of two thirds of OHCs in the mouse organ of Corti [21], while conditional deletion of *Fgfr1* from the cochlear epithelium leads to a loss of almost all OHCs and some IHCs [19,20]. To rule out the effect of genetic background accounting for these differences, we generated *Fgf20* knockout (Fgf20-KO: *Fgf20^-/-^*) and *Fgfr1* conditional knockout (Fgfr1-CKO: *Foxg1^Cre/+^*; *Fgfr1^flox/-^*) mice along with littermate controls (*Fgf20^+/-^*for Fgf20-KO and *Fgfr1^flox/+^*, *Fgfr1^flox/-^*, and *Foxg1^Cre/+^*;*Fgfr1^flox/+^* for Fgfr1-CKO) on a mixed C57BL/6J and 129X1/SvJ genetic background. Fgf20-KO and Fgfr1-CKO mice were generated in separate matings; therefore, some genetic background differences could persist. *Foxg1^Cre^* targets most of the otic vesicle as early as E9.5 [22] and has been used in other studies to conditionally delete *Fgfr1* [18–20]. In the *Fgf20^-^*allele, exon 1 of *Fgf20* is replaced by a sequence encoding a GFP-Cre fusion protein [18]. We also refer to this null allele as *Fgf20^Cre^*.

We examined the cochleae at P0 (Figs 1A and 1B) and quantified the length of the cochlear duct and the total number of IHCs, OHCs, and SCs (Figs 1C-1F), as well as the number of cells along the basal, middle, and apical turns of the cochlear duct (S1A-S1C Figs). Refer to Fig 1G for the positions of basal, middle, and apical turns along the cochlear duct. We identified HCs based on Phalloidin labeling and SCs based on Prox1/Sox2 labeling. IHCs and OHCs were distinguished based on location relative to p75NTR-labeled inner pillar cells (IHCs are neural, or towards the center of the coiled duct; OHCs are abneural).

**Fig 1.**
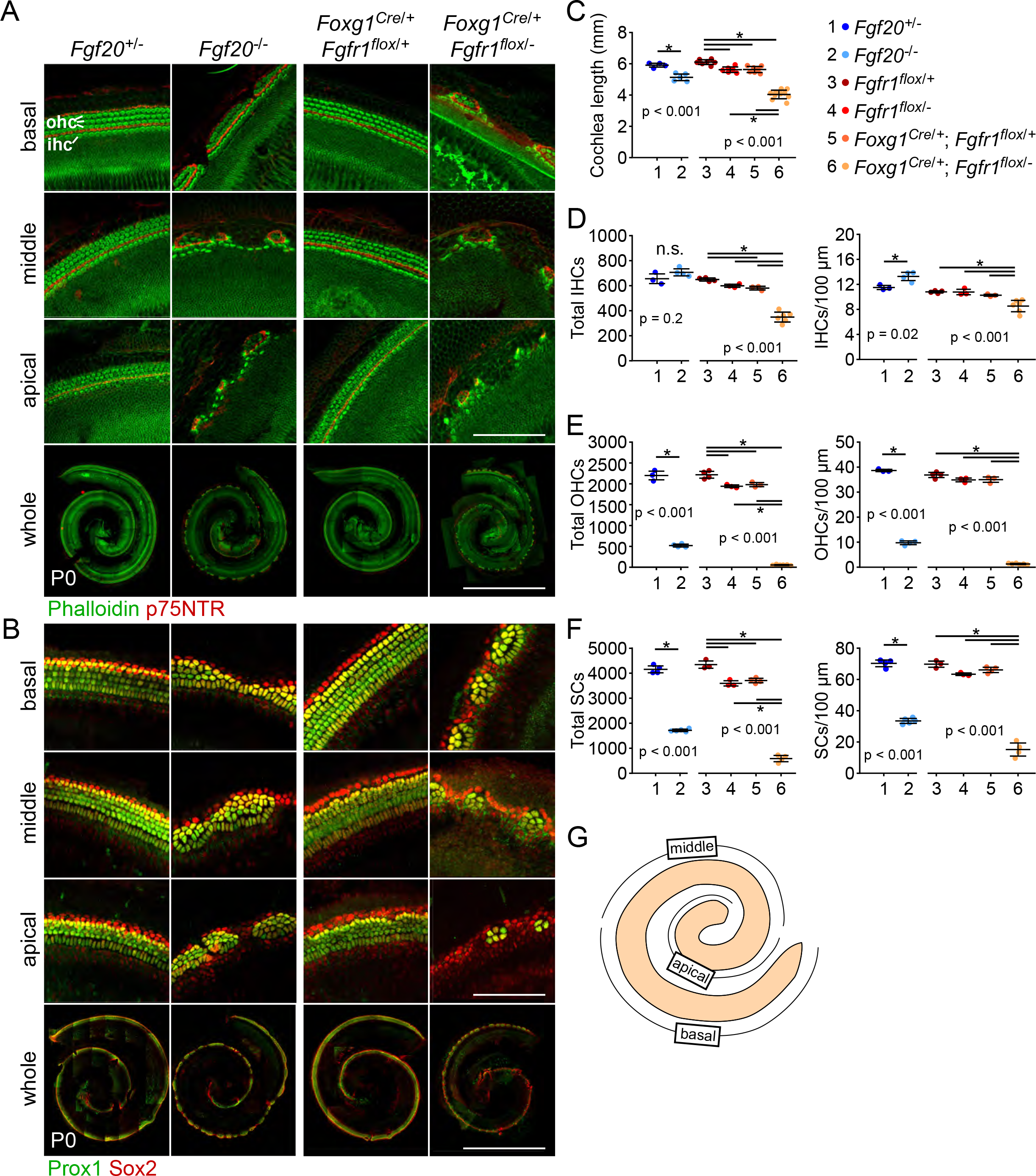
The Fgf20-KO cochlear phenotype is less severe than the Fgfr1-CKO phenotype (A, B) Whole mount cochlea from P0 *Fgf20^+/-^*, *Fgf20^-/-^*, *Foxg1^Cre/+^*;*Fgfr1^flox/+^*, and *Foxg1^Cre/+^*;*Fgfr1^flox/-^* mice showing (A) inner and outer hair cells (IHC and OHC, phalloidin, green) separated by inner pillar cells (p75NTR, red) and (B) supporting cells (Prox1 and Sox2, green/yellow). Magnifications show the basal, middle, and apical turns of the cochlea. Scale bar, 100 µm (magnifications), 1 mm (whole). (C-F) Quantification of (C) cochlear duct length, (D) total IHCs and IHCs per 100 µm of the cochlear duct, (E) total OHCs and OHCs per 100 µm, and (F) total supporting cells (SCs) and SCs per 100 µm at P0. *Fgf20^+/-^* and *Fgf20^-/-^*cochleae results were analyzed by unpaired Student’s t test; *Fgfr1^flox/+^*, *Fgfr1^flox/-^*, *Foxg1^Cre/+^*;*Fgfr1^flox/+^*, and *Foxg1^Cre/+^*;*Fgfr1^flox/-^*cochleae results were analyzed by one-way ANOVA. P values shown are from the t test and ANOVA. * indicates p < 0.05 from Student’s t test or Tukey’s HSD (ANOVA post-hoc); n.s., not significant. Error bars, mean ± SD. (G) Schematic showing the positions of basal, middle, and apical turns along the cochlear duct. See also S1 Fig.

In both Fgf20-KO and Fgfr1-CKO cochleae, there were gaps in the sensory epithelium that lacked HCs and SCs along the entire cochlear duct. Quantitatively, Fgf20-KO cochleae had a 6% reduction in cochlear length compared to control (*Fgf20^+/-^*) cochleae, while Fgfr1-CKO cochleae had a 28% reduction compared to control (*Fgfr1^Cre/+^*;*Fgfr1^flox/+^*). Fgf20-KO did not have a significant reduction in the number of IHCs, while Fgfr1-CKO cochleae had a 40% reduction. Fgf20-KO cochleae only had a 76% reduction in the number of OHCs, while Fgfr1-CKO cochleae had almost a complete lack of OHCs, a 97% reduction. For SCs, Fgf20-KO cochleae had a 59% reduction, while Fgfr1-CKO cochleae had an 84% reduction. These patterns persisted when HC and SC numbers were normalized to cochlear length. These results were all consistent with previous studies [19,21] and showed that the Fgfr1-CKO phenotype is more severe than the Fgf20-KO phenotype in cochlear length and in the number of HCs and SCs. We hypothesize that during organ of Corti development, an additional FGFR1 ligand compensates for the loss of FGF20.

Notably, while the total number of IHCs was decreased in Fgfr1-CKO cochleae, the decrease was only observed in the basal and middle turns of the cochlea, not in the apical turn (S1A Fig). In addition, the number of IHCs normalized to cochlear length was slightly increased in Fgf20-KO cochleae (Fig 1D), and this increase was only prominent in the middle and apical turns of the cochlea, but not in the basal turn (S1A Fig). The increase in IHCs could be explained by the shortened cochlear duct length in Fgf20-KO mice. No such basal/middle/apical turn discrepancies existed in the number of OHCs or SCs in either genotype (S1B and S1C Figs).

Our previous studies also noted that the apical tip of Fgf20-KO cochleae has delayed differentiation relative to control at E16.5 and P0, but catches up by P7 [21]. We confirmed this result, finding that at P0 in control cochleae, sensory epithelium at the apical tip has begun to differentiate, based on phalloidin and p75NTR expression, while in Fgf20-KO cochleae, there was no sign of differentiation at the apical tip. There was a similar delay in differentiation at the apical tip of Fgfr1-CKO cochleae relative to control (S1E Fig). Refer to S1D Fig for the location of the apical tip.

### FGFR1 but not FGF20 regulates *Sox2* expression

Next, we examined *Sox2* expression in Fgf20-KO and Fgfr1-CKO cochleae at E14.5 by RNA in situ hybridization and immunofluorescence. In control cochleae, *Sox2* mRNA and protein were highly expressed in the prosensory domain (Fig 2A, refer to Fig 2C). The expression of *Sox2* was not changed in Fgf20-KO cochleae compared to control; however, it was noticeably decreased in Fgfr1-CKO cochleae (Fig 2A), in agreement with previous findings [19–21]. This indicates that FGFR1 has an additional role, independent of FGF20, in regulating *Sox2*, which is required for prosensory specification [12]. Similar to Sox2, CDKN1B expression in the prosensory domain is also regulated by FGFR1, but not by FGF20 [18,19,21]. We confirmed these results, finding that while CDKN1B expression was not changed in Fgf20-KO cochleae at E14.5 relative to control, it was dramatically downregulated in Fgfr1-CKO cochleae (Fig 2B). This is consistent with the role of Sox2 in regulating CDKN1B expression [12]. We hypothesize that a yet unidentified FGF ligand (in addition to or independent of FGF20) signaling via FGFR1 regulates Sox2 expression (and therefore CDKN1B expression) during prosensory specification, while FGF20 signaling via FGFR1 regulates differentiation (Fig 2D).

**Fig 2.**
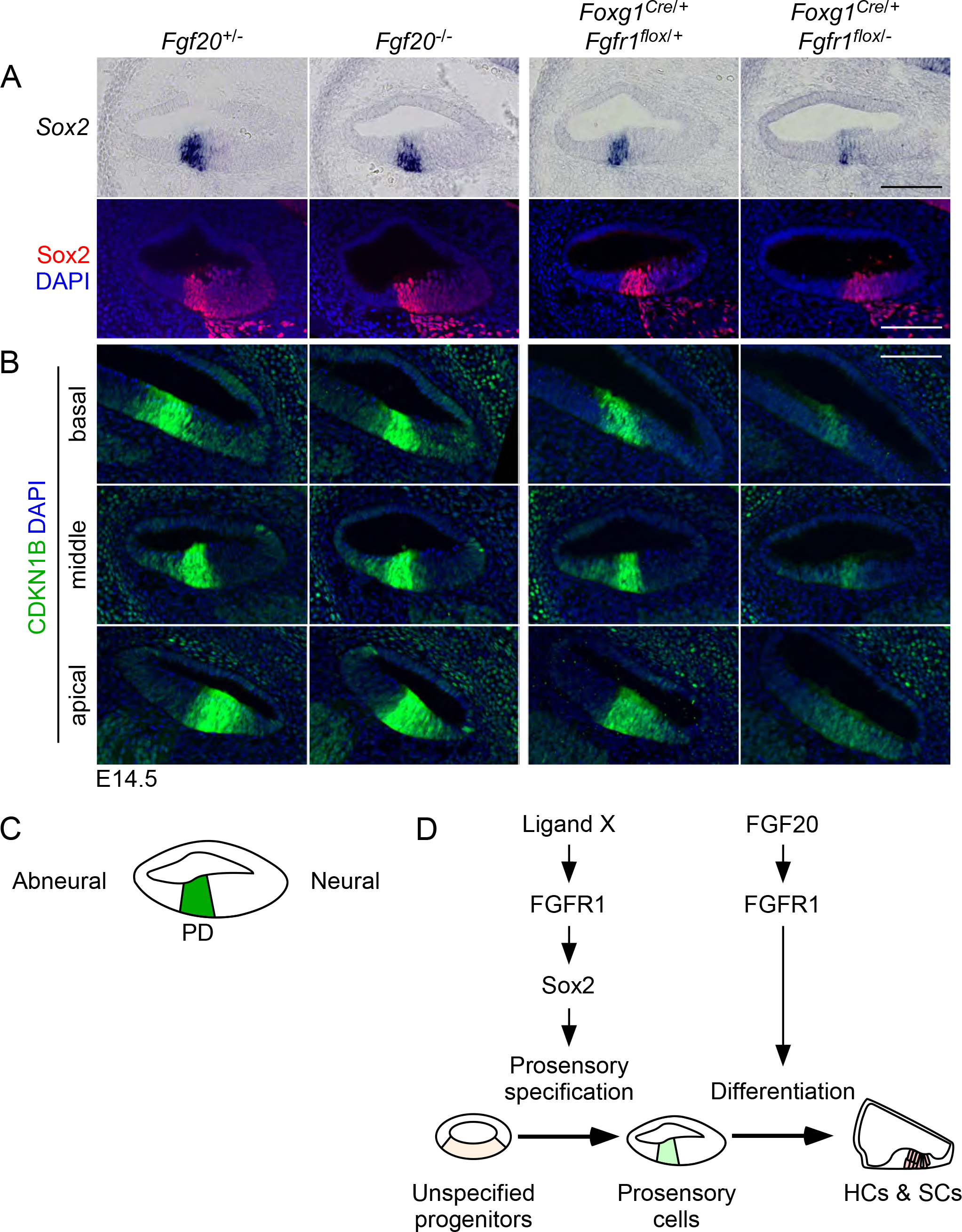
FGFR1 but not FGF20 regulates *Sox2* expression (A) Sections through the middle turn of E14.5 cochlear ducts from *Fgf20^+/-^*, *Fgf20^-/-^*, *Foxg1^Cre/+^*;*Fgfr1^flox/+^*, and *Foxg1^Cre/+^*;*Fgfr1^flox/-^* mice. RNA in situ hybridization (top) and immunofluorescence for *Sox2* (red, bottom), which is expressed in the prosensory domain at this stage. Refer to schematic in (C). (B) Immunofluorescence for CDKN1B (green) in sections through the basal, middle, and apical turns of E14.5 *Fgf20^+/-^*, *Fgf20^-/-^*, *Foxg1^Cre/+^*;*Fgfr1^flox/+^*, and *Foxg1^Cre/+^*;*Fgfr1^flox/-^* cochleae. (C) Schematic of a cross section through the middle turn of the E14.5 cochlear duct, showing the location of the prosensory domain (PD). Neural indicates the side of the duct towards the spiral ganglion cells; abneural indicates away. (D) A model of genetic pathways during organ of Corti development. Ligand X/FGFR1 signaling regulates *Sox2* expression during prosensory specification; FGF20/FGFR1 signaling regulates differentiation. Ligand X may include FGF20, along with another functionally redundant ligand. DAPI, nuclei (blue). Scale bar, 100 µm. See also S2 Fig.

We also wanted to confirm that FGF20 signals to epithelial FGFR1 at around the initiation of differentiation. To do so, we examined the expression of *Etv4* (also known as *Pea3*) and *Etv5* (also known as *Erm*), two well-established downstream effectors of FGF signaling [23], by in situ hybridization. The expression of these two genes are downregulated with FGF signaling inhibition in E14 cochlear explants [17]. At E14.5, there were two domains of *Etv4* and *Etv5* expression in control cochleae: the prosensory domain and the outer sulcus (S2A Fig, brackets). The outer sulcus is the region of the cochlear epithelium abneural to the prosensory domain at E14.5. In Fgf20-KO cochleae, expression of both genes was not detected in the prosensory domain. In Fgfr1-CKO cochleae, expression of both genes was similarly not detected in the prosensory domain. Expression of *Etv4* and *Etv5* in the outer sulcus was not affected in Fgf20-KO and Fgfr1-CKO cochleae (S2A Fig). These results confirm that FGF20 signals through epithelial FGFR1 to the prosensory domain.

Previous studies have also reported a decrease in proliferation in Kölliker’s organ (neural to the prosensory domain, S2B Fig) in Fgfr1-CKO cochleae [20]. We replicated this result by examining EdU (5-ethynyl-2’-deoxyuridine) incorporation at E14.5. *Fgfr1*-CKO mice had a complete lack of EdU-incorporating Kölliker’s organ cells, while Fgf20-KO mice did not show a decrease in EdU incorporation (S2B Fig). This finding is also consistent with an additional FGF ligand signaling via FGFR1, likely at an earlier stage. We do not know whether the proliferation defect in Kölliker’s organ contributes to the reduction in HC and SC number in Fgfr1-CKO mice.

### Genetic rescue of the Fgf20-KO phenotype suggests that FGF20 is required for differentiation

We have previously shown that recombinant FGF9, which is biochemically similar to FGF20 [23,24], is able to rescue the loss of HCs and SCs in Fgf20-KO explant cochleae [21]. Interestingly, while treatment with FGF9 at E13.5 and E14.5 was able to rescue the Fgf20-KO phenotype, treatment at E15.5 was not. This temporal rescue specificity suggests that FGF20 signaling is required for the initiation of HC and SC differentiation.

To confirm the hypothesis that FGF20 is involved in differentiation and not specification (Fig 2D), we sought to more accurately determine the temporal requirement of FGF20 signaling. To achieve this, we developed an *in vivo* genetic rescue model of the Fgf20-KO phenotype by ectopically expressing FGF9. We combined *Fgf20^Cre^* with the *Fgf20^βgal^* [21], *ROSA^rtTA^* [25] and TRE-Fgf9-IRES-eGfp [26] alleles to generate Fgf20-rescue (*Fgf20^Cre/βgal^*;*ROSA^rtTA/+^*;TRE-Fgf9-IRES-eGfp) mice along with littermate controls: Fgf20-het (*Fgf20^Cre/+^*;*ROSA^rtTA/+^*), Fgf9-OA (*Fgf20^Cre/+^*;*ROSA^rtTA/+^*;TRE-Fgf9-IRES-eGfp), and Fgf20-null (*Fgf20^Cre/βgal^*;*ROSA^rtTA/+^*). These mice express the reverse tetracycline transactivator (rtTA) in the *Fgf20^Cre^*lineage, which contains the prosensory domain and Kölliker’s organ at E13.5 to E15.5 [18]. In mice expressing TRE-Fgf9-IRES-eGfp, rtTA drives the expression of FGF9 upon doxycycline (Dox) induction. The *Fgf20^βgal^* allele is another *Fgf20*-null allele, in which exon 1 of *Fgf20* is replaced by a sequence encoding β-galactosidase. We combined *Fgf20^Cre^* with *Fgf20^βgal^*to generate homozygous mutant mice while maintaining a constant dosage of *Fgf20^Cre^*in control and knockouts.

Initially, pregnant dams were fed a Dox diet from E13.5 to E15.5 and pups were harvested at P0 to examine HC and SC development. As expected, Dox treatment itself did not appear to affect HC or SC development in Fgf20-het and Fgf20-null cochleae, both of which showed the expected phenotypes (Figs 3A and 3B). Ectopic expression of FGF9 during these stages also did not affect HC or SC development in Fgf9-OA cochleae, showing that excess FGF20/FGF9 was not sufficient to produce ectopic HCs and SCs. Importantly, ectopic expression of FGF9 resulted in a full rescue of the number and patterning of HCs and SCs in Fgf20-rescue pups. The organ of Corti in these rescue pups had one row of IHCs, three rows of OHCs, and five rows of SCs throughout the entire length of the cochlear duct, without any gaps (Figs 3A and 3B). This shows that FGF20/FGF9 signaling at E13.5-E15.5 is sufficient for HC and SC differentiation. The quantified results from all of the rescue experiments are summarized in Fig 3C, where the number of OHCs and SCs are represented as a percentage of that of Fgf20-het mice treated with the same Dox regimen. All of the quantified data are presented in S3 Fig.

**Fig 3.**
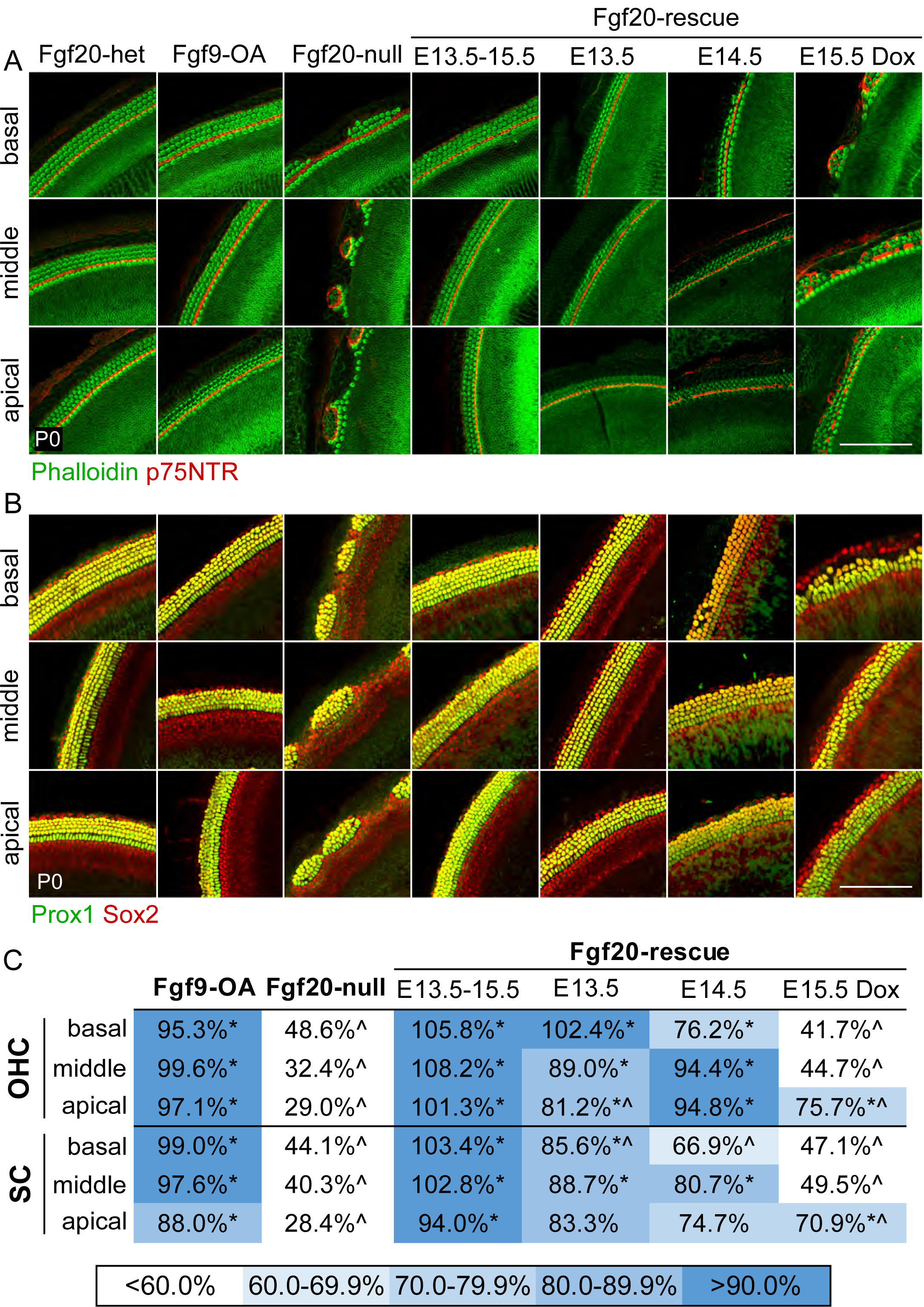
Genetic rescue of the Fgf20-KO phenotype suggests that FGF20 is required for differentiation (A, B) Whole mount cochlea from P0 *Fgf20^+/-^*;*ROSA^rtTA^*(Fgf20-het), *Fgf20^+/-^*;*ROSA^rtTA^*; TRE-Fgf9-IRES-eGfp (Fgf9-OA), *Fgf20^-/-^*;*ROSA^rtTA^*(Fgf20-null), and *Fgf20^-/-^*;*ROSA^rtTA^*; TRE-Fgf9-IRES-eGfp (Fgf20-rescue) mice showing (A) inner and outer hair cells (phalloidin, green) separated by inner pillar cells (p75NTR, red) and (B) supporting cells (Prox1 and Sox2, green/yellow). Images from basal, middle, and apical turns of the cochlea are shown. Fgf20-rescue cochleae from four different doxycycline chow (Dox) regimens are shown (E13.5-E15.5, E13.5, E14.5, and E15.5). Fgf20-het, Fgf9-OA, and Fgf20-null cochleae shown are from the E13.5-E15.5 Dox regimen. Scale bar, 100 µm. (C) Quantification of outer hair cells (OHC) and supporting cells (SC) from P0 Fgf9-OA (Dox regimen: E13.5-E15.5), Fgf20-null (Dox regimen: E13.5-E15.5), and Fgf20-rescue (all four Dox regimens) cochleae, presented as a percentage of the number of cells in Fgf20-het cochleae from the same Dox regimen. * indicates p < 0.05 compared to Fgf20-null cochleae from the same Dox regimen; ^ indicates p < 0.05 compared to Fgf20-het cochleae from the same Dox regimen; Tukey’s HSD (one-way ANOVA post-hoc). See also S3 Fig.

To more precisely determine the timing of rescue sufficiency, we fed pregnant dams Dox for a period of 24 hours starting at E13.5, E14.5, or E15.5. With E13.5 Dox, patterning and OHC number in the basal turn of the cochlea were completely rescued (Fig 3A). However, OHC number in the middle and particularly the apical turns were only partially rescued, resulting in regions with two rows of OHCs instead of three. For instance, in the apical turn, OHC number was restored to 81% of Fgf20-het mice, which is statistically significantly increased compared to Fgf20-null, but also statistically significantly decreased compared to Fgf20-het, indicating partial rescue (Fig 3C). With E14.5 Dox, patterning and OHC number in the middle and apical turns were completely rescued. However, OHC number in the basal turn was not completely rescued, with regions of one or two rows of OHCs, instead of three. With E15.5 Dox, patterning and OHC number was not rescued in the basal and middle turns, as gaps still formed between islands of HCs (Fig 3A). However, OHC number in the apical turn was partially rescued, with two or three rows of OHCs not separated by gaps towards the tip of the apex. In all of these experiments, the rescue of SCs followed the same pattern as that of OHCs (Fig 3B).

These rescue results show that FGF20/FGF9 is sufficient for OHC and SC differentiation in the basal turn of the cochlea at E13.5, in the middle and apical turns at E14.5-E15.5, and in the tip of the apical turn at E15.5. Since the initiation of HC and SC differentiation occurs in the base/mid-base of the cochlea at E13.5 and progresses apically over the next few days, these results strongly imply that FGF20 functions during the initiation of differentiation, rather than prosensory specification, consistent with our model (Fig 2D).

### Decrease in *Sox2* expression results in similar phenotypes as disruptions to FGFR1 signaling

Our results and previous findings suggest that FGFR1 regulates prosensory specification via Sox2 [19]. Mice with an inner ear-specific *Sox2* hypomorphic mutation (*Sox2^Ysb/Ysb^*, see below) have defects in prosensory specification, accounting for a small loss of HCs and SCs, whereas mice with inner-ear specific *Sox2* null mutations have a complete lack of prosensory specification and a complete absence of sensory epithelium [12]. To examine how much the reduction in *Sox2* expression in Fgfr1-CKO cochlea contributes to the phenotype at P0, we combined the *Sox2^-^* (*Sox2* constitutive null) and *Sox2^Ysb^* alleles to closely examine the effects of reduction in *Sox2* expression on organ of Corti development, on a similar genetic background as our Fgf20-KO and Fgfr1-CKO mice. We hypothesized that if *Fgfr1* acts upstream of *Sox2*, then reducing *Sox2* expression should at least partially recapitulate the Fgfr1-CKO cochlea phenotype. The *Sox2^Ysb^*allele is a regulatory mutant in which transgene insertion in chromosome 3 disrupts some otic enhancers, resulting in hypomorphic *Sox2* expression in the inner ear [11,12].

We generated a *Sox2* allelic series of mice with the following genotypes, in order of highest to lowest levels of *Sox2* expression: *Sox2^+/+^*(wildtype), *Sox2^Ysb/+^*, *Sox2^Ysb/Ysb^*, and *Sox2^Ysb/-^*. In this allelic series, decrease in *Sox2* expression had a dose-dependent effect on cochlea length at P0 (Figs 4A-4C). *Sox2^Ysb/+^* cochleae had a 6% reduction in length compared to wildtype (although not statistically significant), *Sox2^Ysb/Ysb^* cochleae had a 24% reduction, and *Sox2^Ysb/-^*had a 46% reduction. *Sox2^Ysb/+^* organ of Corti developed relatively normally, with three rows of OHCs and one row of IHCs (Fig 4A). Interestingly, there were occasional ectopic IHCs medial (neural) to the normal row of IHCs, especially in the middle and apical turns of the *Sox2^Ysb/+^*cochlea (Fig 4A, arrowheads). However, there was no significant increase in IHC number (total or normalized to length) compared to wildtype cochleae (Fig 4D). The *Sox2^Ysb/Ysb^*cochlea appeared much more abnormal, with gaps in the sensory epithelium that lacked HCs and SCs in the basal turn (Figs 4A and 4B), similar to what was observed previously [12]. Moreover, at the base, in the sensory islands between the gaps, there were often four rows of OHCs and six rows of SCs. In the middle and apical turns, there were the normal three rows of OHCs and five rows of SCs. There were also numerous ectopic IHCs throughout the middle and apical turns, sometimes forming an entire second row of cells (Fig 4A), resulting in increased number of IHCs in the middle turn compared to wildtype (S4A Fig). However, the total and length-normalized number of IHCs in *Sox2^Ysb/Ysb^* cochleae did not significantly differ from that of wildtype cochleae (Fig 4D). In terms of OHCs, *Sox2^Ysb/Ysb^*cochleae exhibited a 40% decrease in total number compared to wildtype cochleae (Fig 4E). This decrease was not quite as severe when normalized to cochlear length (21% decrease). Strikingly, *Sox2^Ysb/-^* cochleae lacked almost all HCs and SCs, except in the apical turn (Figs 4A and 4B). The decrease in OHC number (93%) in *Sox2^Ysb/-^* cochleae compared to wildtype was more severe than the decrease in IHC number (75%). Notably, IHC number was significantly decreased in the basal and middle turns, but not in the apical turn (S4A Fig). OHC number was significantly decreased throughout all three turns (S4B Fig). In all of these genotypes, the number of SCs followed the pattern of loss of OHCs (Fig 4F and S4C Fig). Interestingly, while *Sox2^Ysb/-^* cochleae almost completely lacked HCs and SCs in the basal and middle turns, in 7 of 11 *Sox2^Ysb/-^*cochleae examined, one or two small islands of HCs or SCs were found at the basal tip (S4D Fig).

**Fig 4.**
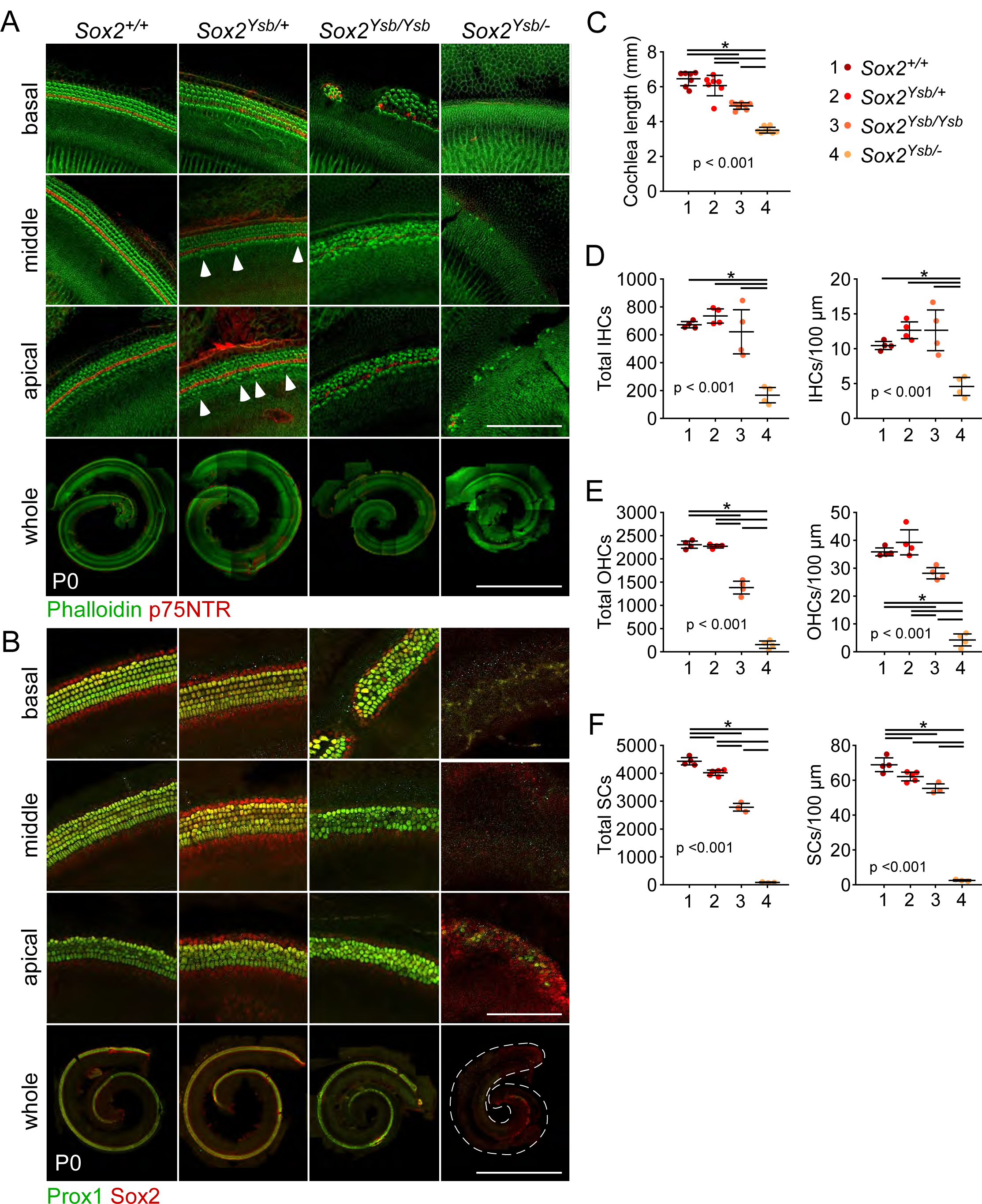
Decrease in Sox2 expression results in similar phenotypes as disruptions to FGFR1 signaling (A, B) Whole mount cochlea from P0 mice from the *Sox2* allelic series (in order of highest to lowest levels of *Sox2* expression: *Sox2^+/+^*, *Sox2^Ysb/+^*, *Sox2^Ysb/Ysb^*, and *Sox2^Ysb/-^*) showing inner and outer hair cells (phalloidin, green) separated by inner pillar cells (p75NTR, red) and (B) supporting cells (Prox1 and Sox2, green/yellow). Magnifications show the basal, middle, and apical turns of the cochlea. Arrowheads indicate ectopic inner hair cells. Scale bar, 100 µm (magnifications), 1 mm (whole). (C-F) Quantification of (C) cochlear duct length, (D) total inner hair cells (IHCs) and IHCs per 100 µm of the cochlear duct, (E) total outer hair cells (OHCs) and OHCs per 100 µm, and (F) total supporting cells (SCs) and SCs per 100 µm at P0. P values shown are from one-way ANOVA. * indicates p < 0.05 from Tukey’s HSD (ANOVA post-hoc). Error bars, mean ± SD. See also S4 Fig.

Overall, these results showed that the basal end of the cochlea is more sensitive to the loss of *Sox2* expression than the apical end. Furthermore, while both IHCs and OHCs were affected, OHCs were more sensitive to decrease in *Sox2* expression than IHCs. Importantly, both of these features were found in Fgfr1-CKO cochleae, where the decrease in IHCs was only found in the basal and middle turns and there were almost no OHCs along the entire cochlear duct (S1A and S1B Fig). Therefore, we conclude that decrease in *Sox2* expression, leading to defects in prosensory specification, could account for the Fgfr1-CKO phenotype. Furthermore, the decrease in *Sox2* expression could also account for the difference in severity between the Fgf20-KO and Fgfr1-CKO phenotypes, since Fgf20-KO cochleae, which had normal *Sox2* expression, did not have a decrease in the number of IHCs, unlike Fgfr1-CKO and *Sox2^Ysb/-^* cochleae.

### Decrease in levels of Sox2 expression delays prosensory specification

We sought to determine why a decrease in *Sox2* expression more severely affected the basal end of the cochlear duct. Initially, we examined *Sox2* expression at E14.5. As expected, Sox2 expression was almost completely absent in *Sox2^Ysb/-^* cochleae (S5A Fig). This decrease in expression was not more severe at the basal turn of the cochlear duct, relative to the middle and apical turns, suggesting that the more severe basal phenotype in *Sox2^Ysb/-^*cochleae cannot be explained by differential Sox2 expression. Similarly, CDKN1B expression was downregulated in the prosensory domain of *Sox2^Ysb/-^*cochleae, consistent with previous studies [12]. Interestingly, the decrease in expression was also not more severe at the basal turn relative to the middle and apical turns (S5B Fig). Using CDKN1B as a marker of prosensory specification, this suggests that the more severe basal phenotype also cannot be explained by differential regulation of prosensory specification along the length of the cochlea.

As described in the introduction, waves of cell cycle exit (marking the completion of prosensory specification) and initiation of differentiation travel in opposite directions along the cochlear duct during development, resulting in the basal end of the cochlear duct differentiating immediately after specification. The apical end, meanwhile, exhibits a delay in differentiation, resulting in a longer temporal buffer between specification and differentiation. In this developmental pathway, specification must be completed prior to the initiation of differentiation. We reasoned, therefore, that disruptions to the timing of prosensory specification will preferentially interfere with basal sensory epithelia development, potentially accounting for the more severe basal phenotype in *Sox2* hypomorphs.

To test this hypothesis, we examined cell cycle exit in the prosensory domain via Ki67 expression, as a marker of the status of prosensory specification. Ki67 is expressed by cycling cells, but not cells in G_0_ [27]. In the developing cochlea at around E12.5 to E15.5, cells of the prosensory domain, sometimes referred to as the zone of non-proliferation, have turned off or are beginning to turn off Ki67 [3]. At E14.5 in *Sox2^Ysb/+^* cochleae, the prosensory domain along most of the cochlear duct (serial sections 2-6) has turned off Ki67 expression, except at the very base (serial section 1; Fig 5A, brackets). See graphical summary below Fig 5A; also see S5C Fig for serial “mid-modiolar” sections through the cochlea. This indicates that the wave of cell cycle exit, which starts at the apex, has reached the very base of the cochlear duct. However, in *Sox2^Ysb/-^* cochleae, only the prosensory domain at the apical turn of the cochlear duct (serial section 6) has turned off Ki67, not at the mid-basal or basal turns (serial sections 1-3); the middle turns (serial sections 4 and 5), meanwhile, were just starting to turn off Ki67 (Fig 5A, brackets). This indicates that cell cycle exit has only reached the middle turn, suggesting a delay in prosensory specification. In addition, the nuclei of prosensory domain cells shift away from the luminal surface of the cochlear epithelium upon specification [28]. This basal shift of nuclei localization within the cell leaves a blank space between DAPI-stained nuclei and the luminal surface of the cochlear duct, which can be visualized in all six serial sections in *Sox2^Ysb/+^*cochleae at E14.5 (Fig 5A, asterisks). However, in *Sox2^Ysb/-^*cochleae, cells of the prosensory domain mostly did not exhibit this nuclei shift at E14.5.

**Fig 5.**
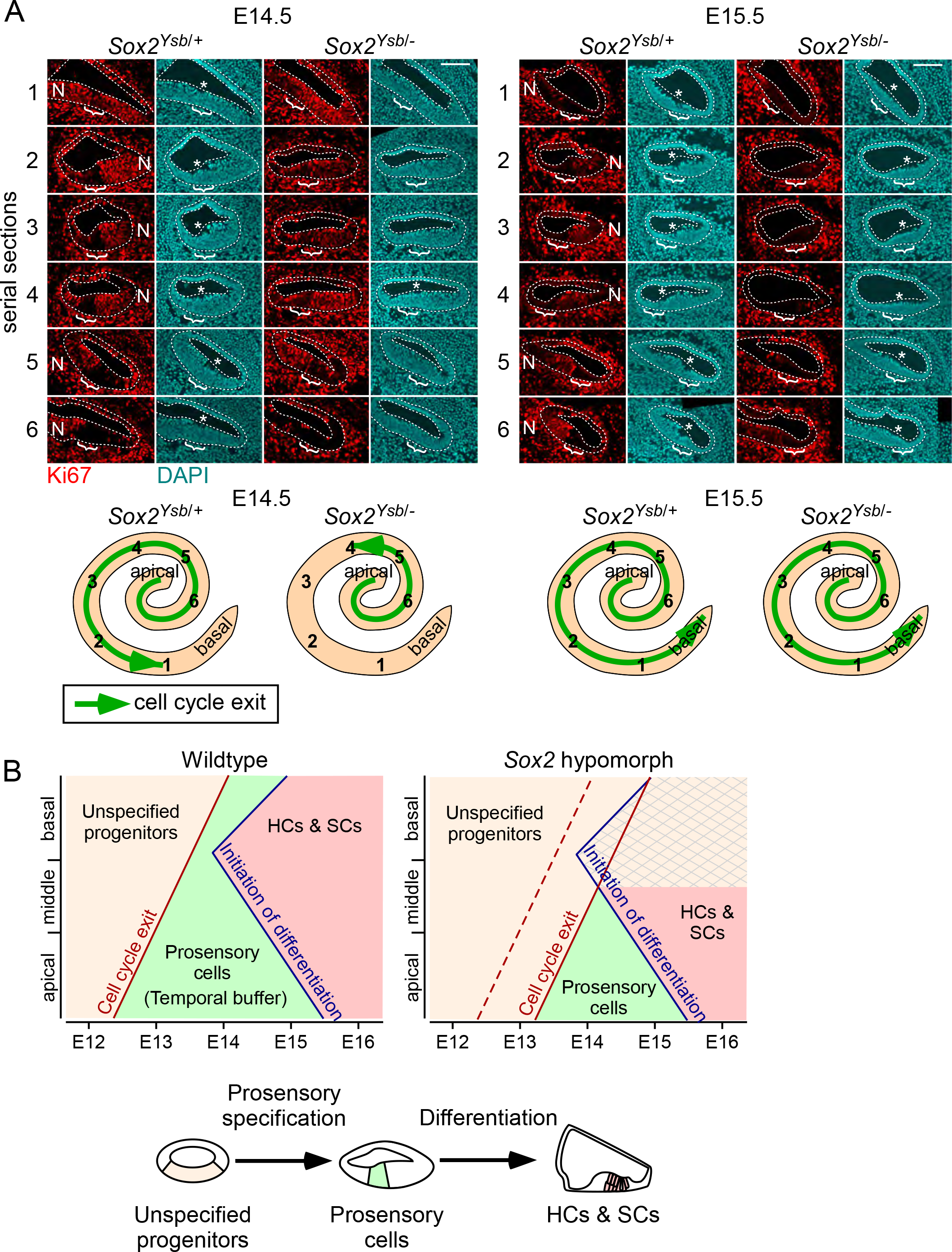
Decrease in levels of Sox2 expression delays prosensory specification (A) Serial sections (1-6) through the duct of E14.5 and E15.5 *Sox2^Ysb/+^* and *Sox2^Ysb/-^* cochleae. Immunofluorescence for Ki67 (red) and DAPI (nuclei, cyan). Cochlear epithelium is outlined. Bracket indicates prosensory domain. * indicates shift of prosensory nuclei away from the luminal surface of the epithelium. N, neural side. Scale bar, 100 µm. Whole mount cochlear duct schematics show relative positions of the serial sections and progression of cell cycle exit (green arrow). (B) A model of organ of Corti development showing embryonic staging (x-axis) and location along the cochlear duct (basal, middle, and apical turns, y-axis). Development occurs in two stages: unspecified progenitors (tan shading) undergo specification and cell cycle exit to become prosensory cells (green shading), which then differentiate into hair cells and supporting cells (HCs & SCs; red shading). In wildtype cochleae, cell cycle exit (indicating completion of specification) begins at the apex of the cochlea and proceeds basally. Afterwards, differentiation initiates at the mid-base of the cochlea and proceeds basally and apically. Temporal buffer (green shading) refers to the time between cell cycle exit and initiation of differentiation. In *Sox2* hypomorph cochleae, specification and cell cycle exit are delayed, resulting in failure to complete specification before initiation of differentiation towards the basal end of the cochlea (crosshatch pattern). See also S5 Fig.

At E15.5, the prosensory domain along the entire length of the cochlear duct has turned off Ki67 expression in both *Sox2^Ysb/+^*and *Sox2^Ysb/-^* cochleae (Fig 5A, brackets), indicating that prosensory specification in *Sox2^Ysb/-^*cochleae has caught up by this stage. Prosensory nuclei localization has also begun to catch up at E15.5 in *Sox2^Ysb/-^* cochleae (Fig 5A, asterisks). Overall, these results suggest that prosensory specification is delayed in *Sox2^Ysb/-^* cochleae, but not permanently disrupted.

By definition, prosensory specification must occur prior to differentiation to generate HCs and SCs. Therefore, the period of time in between cell cycle exit and the initiation of differentiation represents a temporal buffer (Fig 5B, green shading) preventing differentiation from initiating prior to specification. As differentiation begins in the basal/mid-basal cochlear turns shortly after specification, the delay in specification in *Sox2^Ysb/-^*cochleae leads to progenitors not having been specified in time for differentiation at the basal end of the cochlear duct (Fig 5B, crosshatch pattern). We propose that this at least partially explains why the basal end of the cochlea is more sensitive to decreases in the level of *Sox2* expression. Moreover, since differentiation begins in the mid-base and spreads to the rest of the base and apex, progenitors at the basal tip in *Sox2^Ysb/-^* cochleae may still undergo specification prior to differentiation. This may explain why small islands of HCs and SCs are sometimes seen in the basal tip of *Sox2^Ysb/-^*cochleae (S4D Fig).

### *Sox2* is upstream of *Fgf20*

While the delay in prosensory specification can explain the preferential loss of sensory epithelium from the basal end of *Sox2* hypomorph cochleae, it does not readily explain the preferential loss of OHCs, relative to IHCs. Since this preference for OHC loss is reminiscent of the *Fgf20*/*Fgfr1* deletion phenotypes, we investigated the possibility that *Sox2* may be upstream of FGF20-FGFR1 signaling. Interestingly, both *Etv4* and *Etv5* were dramatically downregulated in the prosensory domain of *Sox2^Ysb/-^* cochleae compared to control (Fig 6A). This shows that FGF20-FGFR1 signaling was disrupted in the *Sox2* hypomorph cochleae. Examination of *Fgfr1* and *Fgf20* expression by in situ hybridization revealed that while *Fgfr1* expression did not appear to be affected in *Sox2^Ysb/-^* cochleae at E14.5, *Fgf20* expression was absent (Fig 6B). This suggests that while *Fgfr1* functions upstream of *Sox2*, *Fgf20* is downstream of *Sox2*. This model predicts that *Fgf20* expression would be downregulated in Fgfr1-CKO cochleae, which was confirmed by in situ hybridization (Fig 6C).

**Fig 6.**
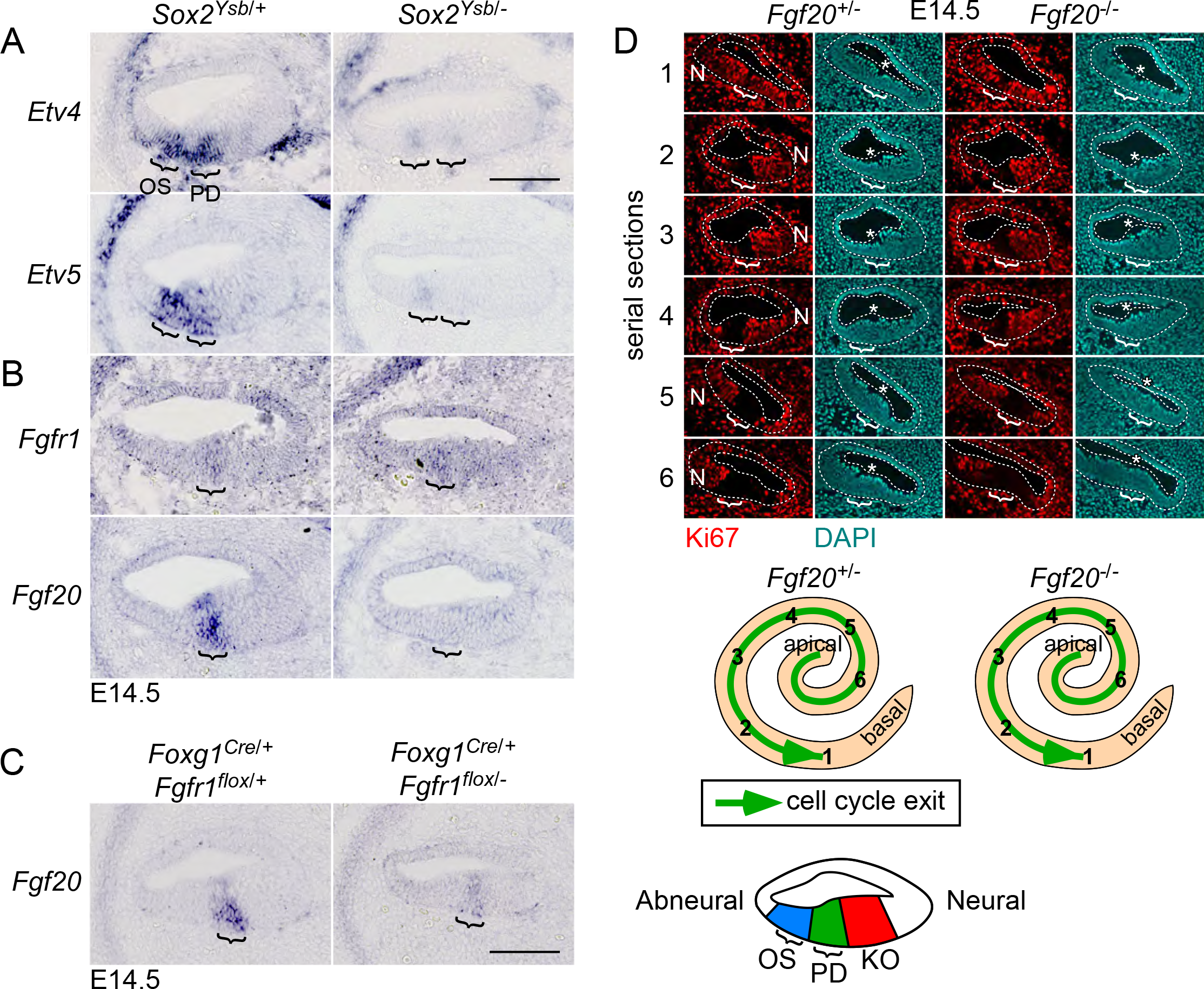
*Sox2* is upstream of *Fgf20* (A-C) Sections through the middle turn of E14.5 cochleae. (A) RNA in situ hybridization for *Etv4* and *Etv5* in *Sox2^Ysb/+^* and *Sox2^Ysb/-^* cochleae. The two brackets indicate *Etv4/5* expression in the outer sulcus (OS, left) and prosensory domain (PD, right). Refer to schematic at the bottom right of the figure. (B) RNA in situ hybridization for *Fgfr1* and *Fgf20* in *Sox2^Ysb/+^* and *Sox2^Ysb/-^* cochleae. Bracket indicates *Fgfr1*/*Fgf20* expression in the prosensory domain. (C) RNA in situ hybridization for *Fgf20* in *Foxg1^Cre/+^*;*Fgfr1^flox/+^* and *Foxg1^Cre/+^*;*Fgfr1^flox/-^*cochleae. Bracket indicates *Fgf20* expression in the prosensory domain. (D) Serial sections (1-6) through the duct of E14.5 *Fgf20^+/-^* and *Fgf20^-/-^* cochleae. Immunofluorescence for Ki67 (red) and DAPI (nuclei, cyan). Cochlear epithelium is outlined. Bracket indicates prosensory domain. * indicates shift of prosensory nuclei away from the luminal surface of the epithelium. N, neural side. Scale bar, 100 µm. Whole mount cochlear duct schematics show relative positions of the serial sections and progression of cell cycle exit (green arrow). OS, outer sulcus; PD, prosensory domain; KO, Kölliker’s organ. Scale bar, 100 µm. See also S6 Fig.

The above results indicate that the loss of *Fgf20* could partially account for the *Sox2^Ysb/-^* phenotype. To determine whether loss of *Fgf20* also causes delayed prosensory specification, we examined Ki67 expression in Fgf20-KO cochleae. At E14.5, there was no detectable delay in cell cycle exit in Fgf20-KO cochleae, as loss of Ki67 expression reached the base (serial section 1) in both control and Fgf20-KO cochleae (Fig 6D, brackets). See S6A Fig for serial “mid-modiolar” sections through the cochlea. There was also no detectable delay in prosensory basal nuclei shift in Fgf20-KO cochleae (Fig 6D, asterisks). These results were expected as the Fgf20-KO phenotype is not more severe at the basal end of the cochlear duct. This is also consistent with *Fgf20* being required during differentiation rather than prosensory specification (Fig 2D). However, these results do not answer whether and how the loss of *Fgf20* contributes to the *Sox2* hypomorph phenotype.

We also asked whether decrease in *Sox2* expression can account for the absence of proliferation in Kölliker’s organ of Fgfr1-CKO cochleae. Interestingly, EdU-incorporation was decreased in Kölliker’s organ in *Sox2^Ysb/-^* cochleae at E14.5, especially in the region adjacent to the prosensory domain (S6B Fig, bracket). However, EdU-incorporation was not completely absent from Kölliker’s organ, unlike in Fgfr1-CKO cochleae. This suggests that loss of *Sox2* in combination with other factors contributes to Kölliker’s organ phenotype in Fgfr1-CKO cochleae.

### *Sox2* and *Fgf20* interact during cochlea development

To explore how the loss of *Fgf20* contributes to the *Sox2* hypomorph phenotype, we combined the *Fgf20^-^* and *Sox2^Ysb^* alleles to generate *Fgf20* and *Sox2* compound mutants. We also hypothesized that reducing *Sox2* expression in Fgf20-KO mice would recapitulate (or phenocopy) the more severe Fgfr1-CKO phenotype. We interbred F1 mice from the same parents to generate nine different F2 genotypes encompassing all possible combinations of the *Fgf20^-^* and *Sox2^Ysb^*alleles: *Fgf20^+/+^*;*Sox2^+/+^*, *Fgf20^+/+^*;*Sox2^Ysb/+^*, *Fgf20^+/-^*;*Sox2^+/+^*, *Fgf20^+/-^*;*Sox2^Ysb/+^*, *Fgf20^+/+^*;*Sox2^Ysb/Ysb^*, *Fgf20^+/-^*;*Sox2^Ysb/Ysb^*, *Fgf20^-/-^*;*Sox2^+/+^*, *Fgf20^-/-^*;*Sox2^Ysb/+^*, and *Fgf20^-/-^*;*Sox2^Ysb/Ysb^*(Figs 7A and 7B and data not shown). At P0, an overview of HCs and SCs showed that the *Fgf20^+/-^*;*Sox2^Ysb/+^* phenotype mostly resembled that of *Fgf20^+/+^*;*Sox2^+/+^*, *Fgf20^+/+^*;*Sox2^Ysb/+^*, and *Fgf20^+/-^*;*Sox2^+/+^* cochleae, except for the prevalence of ectopic IHCs (Fig 7A, arrowheads). The *Fgf20^+/-^*;*Sox2^Ysb/Ysb^*phenotype mostly resembled that of *Fgf20^+/+^*;*Sox2^Ysb/Ysb^*cochleae, but with more gaps in the basal cochlear turn and two rows of IHCs throughout the length of the cochlear duct, except where there were gaps. The *Fgf20^-^/^-^*;*Sox2^Ysb/+^*phenotype mostly resembled that of *Fgf20^-/-^*;*Sox2^+/+^*cochleae, but with smaller sensory islands in between gaps. The *Fgf20^-/-^*;*Sox2^Ysb/Ysb^*phenotype appeared by far the most severe, with almost a complete absence of IHCs, OHCs, and SCs from the basal turn, and tiny sensory islands in the middle turn; however, the apical turn appeared similar to that of *Fgf20^-^/-*;*Sox2^Ysb/+^* and *Fgf20^-/-^*;*Sox2^+/+^*cochleae (Figs 7A and 7B).

**Fig 7.**
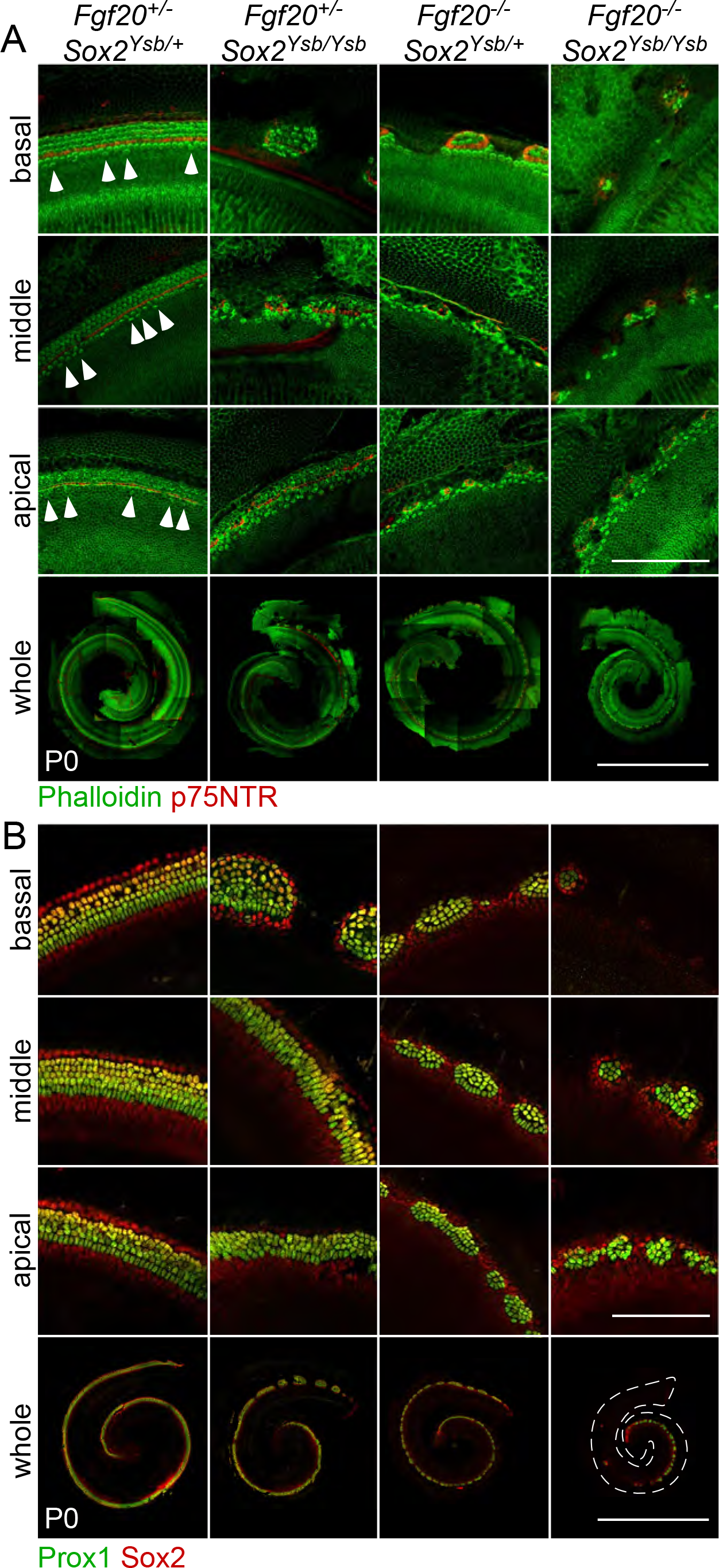
*Sox2* and *Fgf20* interact during cochlea development (A, B) Whole mount cochlea from P0 *Fgf20^+/-^*;*Sox2^Ysb/+^*, *Fgf20^+/-^*;*Sox2^Ysb/Ysb^*, *Fgf20^-/-^*;*Sox2^Ysb/+^*, and *Fgf20^-/-^*;*Sox2^Ysb/Ysb^* mice showing (A) inner and outer hair cells (phalloidin, green) separated by inner pillar cells (p75NTR, red) and (B) supporting cells (Prox1 and Sox2, green/yellow). Magnifications show the basal, middle, and apical turns of the cochlea. Arrowheads indicate ectopic inner hair cells. Scale bar, 100 µm (magnifications), 1 mm (whole).

Quantification of the phenotypes are presented in Figs 8B-8E and S7B-S7D Figs. We analyzed the quantified P0 phenotype via two-way ANOVA with the two factors being gene dosage of *Fgf20* (levels: *Fgf20^+/+^*, *Fgf20^+/-^*, *Fgf20^-/-^*) and *Sox2* (levels: *Sox2^+/+^*, *Sox2^Ysb/+^*, *Sox2^Ysb/Ysb^*). Results from the two-way ANOVA and post-hoc Tukey’s HSD are presented in Figs 8A, 8F, and S7A and S8 Figs. Cochlear length and the total number of IHCs, OHCs, and SCs were all significantly affected by both the *Fgf20* dosage and the *Sox2* dosage, as well as an interaction between the two factors (Figs 8A-8E). The statistically significant interaction between *Fgf20* and *Sox2* dosages suggests that *Fgf20* and *Sox2* have a genetic interaction in regulating cochlear length as well as the number of IHCs, OHCs, and SCs (Fig 8A). Notably, *Fgf20^+/-^*;*Sox2^Ysb/Ysb^*cochleae had significantly fewer OHCs and SCs than *Fgf20^+/+^*;*Sox2^Ysb/Ysb^*cochleae, and *Fgf20^-/-^*; *Sox2^Ysb/+^* cochleae had significantly fewer OHCs than *Fgf20^-/-^*;*Sox2^+/+^*cochleae (Fig 8F). Importantly, *Fgf20^-/-^*;*Sox2^Ysb/Ysb^* cochleae had decreased total and length-normalized number of IHCs, which was not observed in any of the other genotypes, strongly supporting a genetic interaction between *Fgf20* and *Sox2* (*Fgf20^+/+^*;*Sox2^Ysb/Ysb^*cochleae did have a slight decrease in the total number IHCs, but not in the length-normalized number of IHCs).

**Fig 8.**
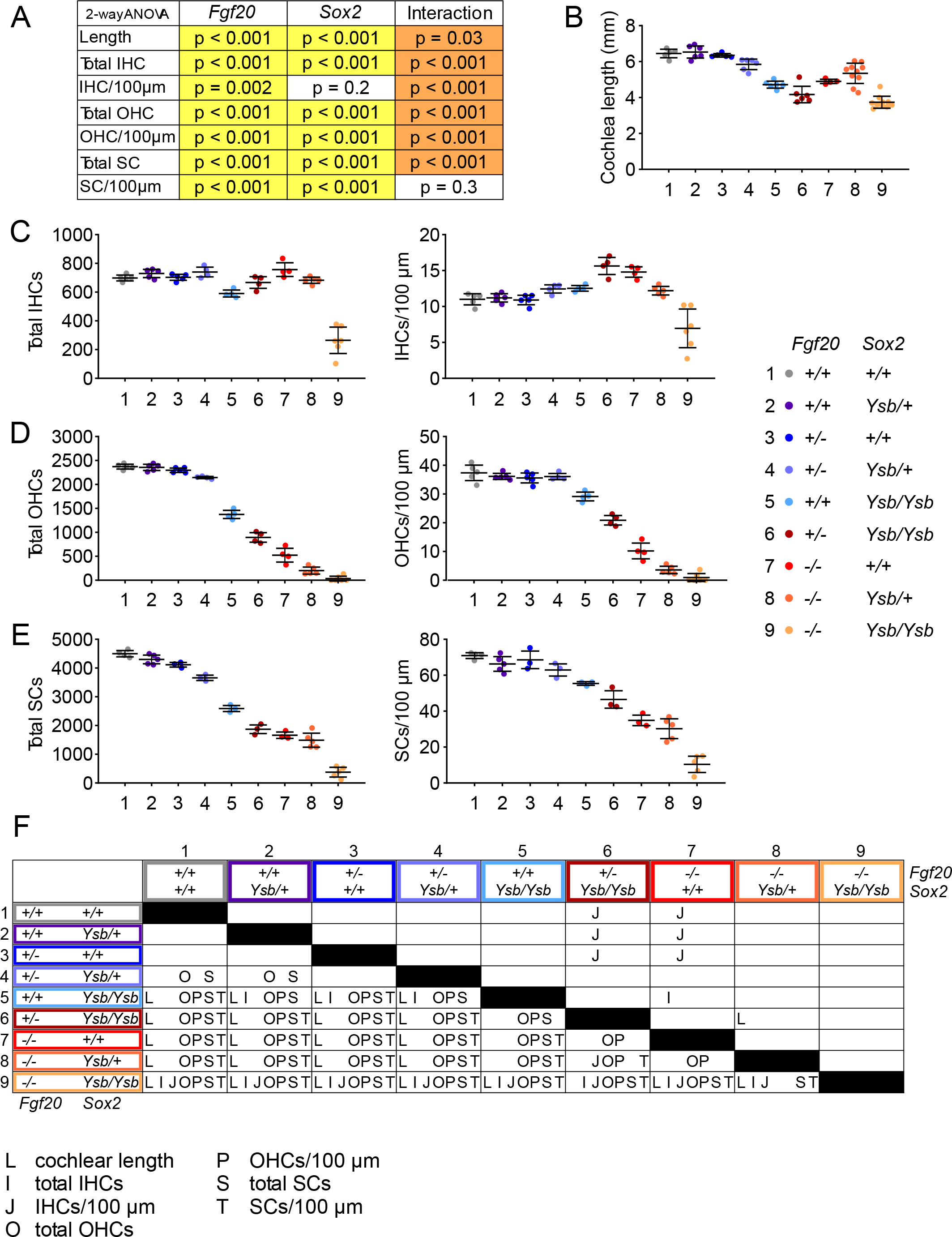
*Sox2* and *Fgf20* interact during cochlea development—quantitative analysis (A) P values from two-way ANOVA analyzing the quantification results in (B-E). The two factors analyzed are *Fgf20* (*Fgf20^+/+^*, *Fgf20^+/-^*, *Fgf20^-/-^*) and *Sox2* (*Sox2^+/+^*, *Sox2^Ysb/+^*, *Sox2^Ysb/Ysb^*) gene dosage. A p value < 0.05 (yellow highlight) for *Fgf20* or *Sox2* indicates that the particular factor (independent variable) has a statistically significant effect on the measurement (dependent variable). Whereas a p value < 0.05 (orange highlight) for Interaction indicates a statistically significant interaction between the effects of the two factors on the measurement. (B-E) Quantification of (B) cochlear duct length, (C) total inner hair cells (IHCs) and IHCs per 100 µm of the cochlear duct, (D) total outer hair cells (OHCs) and OHCs per 100 µm, and (E) total supporting cells (SCs) and SCs per 100 µm at P0 in *Fgf20^+/+^*;*Sox2^+/+^*, *Fgf20^+/+^*;*Sox2^Ysb/+^*, *Fgf20^+/-^*;*Sox2^+/+^*, *Fgf20^+/-^*;*Sox2^Ysb/+^*, *Fgf20^+/+^*;*Sox2^Ysb/Ysb^*, *Fgf20^+/-^*;*Sox2^Ysb/Ysb^*, *Fgf20^-/-^*;*Sox2^+/+^*, *Fgf20^-/-^*;*Sox2^Ysb/+^*, and *Fgf20^-/-^*;*Sox2^Ysb/Ysb^*cochleae. Error bars, mean ± SD. (F) Results from post-hoc Tukey’s HSD analyzing the quantification results in (B-E). Letters (L, I, J, O, P, S, T; representing each measurement in panels B-E) indicate a statistically significant decrease (p < 0.05) when comparing the row genotype against the column genotype. L, cochlear length; I, total IHCs; J, IHCs/100 µm; O, total OHCs; P, OHCs/100 µm; S, total SCs; T, SCs/100 µm. See also S7 and S8 Figs.

Interestingly, while the total number of IHCs was decreased in *Fgf20^-/-^*;*Sox2^Ysb/Ysb^* cochleae relative to all other genotypes, this decrease was only found in the basal and middle turns, but not the apical turn (S7B and S8 Figs). No such basal/middle/apical turn discrepancies existed in the number of OHCs or SCs (S7C, S7D, and S8 Figs). This is reminiscent of the Fgfr1-CKO and *Sox2^Ysb/-^* phenotypes.

To ensure that the *Fgf20^-^* and *Sox2^Ysb^*interaction is not purely an artifact of the *Sox2^Ysb^* allele, we generated *Fgf20^+/+^*;*Sox2^+/+^*(wildtype), *Fgf20^+/-^*;*Sox2^+/+^* (Fgf20-het), *Fgf20^+/+^*;*Sox2^+/-^* (Sox2-het), and *Fgf20^+/-^*;*Sox2^+/-^*(double het) mice to look for an interaction between the *Fgf20^-^* and *Sox2^-^* alleles (S9A Fig). At P0, cochlear length did not significantly differ among the four genotypes (S9B Fig). HC quantification showed that neither *Fgf20* nor *Sox2* exhibited haploinsufficiency for total or length-normalized number of IHCs or OHCs (S9C and S9D Figs). However, in Fgf20-het and much more so in Sox2-het cochleae, occasional ectopic IHCs can be found in the middle and apical turns of the cochlear duct (S9A Fig, arrowheads). Interestingly, in double het cochleae, many more ectopic IHCs were found, even in the basal turn. These ectopic IHCs led to an increase in the total and length-normalized number of IHCs in double het cochleae, compared to wildtype (S9C Fig). Notably, a significant increase in IHCs was only found in the basal turn, not the middle or apical turns (S9E Fig). In the basal turn, IHC number was significantly increased in double het cochleae compared to wildtype, Fgf20-het, and Sox2-het cochleae. Double het cochleae also had a significant decrease in total and length-normalized number of OHCs compared to wildtype (S9D Fig). Again, a significant decrease in OHCs was only found in the basal turn, not the middle or apical turns (S9F Fig). These results confirm a genetic interaction between *Fgf20* and *Sox2*.

### Loss of *Fgf20* does not further delay prosensory specification in *Sox2* **hypomorph cochleae**

We propose that the *Fgf20^-/-^*;*Sox2^Ysb/Ysb^*phenotype lies in between that of Fgfr1-CKO and *Sox2^Ysb/-^* in terms of severity of reductions in cochlear length and in the number of HCs and SCs. We further hypothesize that these three phenotypes form a continuum with the Fgf20-KO phenotype (Fig 9A). Along this continuum, all four genotypes lack FGF20 signaling, but vary in the level of *Sox2* expression and phenotype severity in the basal end of the cochlear duct and the outer compartment (outer rows of OHCs and SCs). From this, and from the *Fgf20*^-^ and *Sox2^Ysb^*series of alleles, we conclude that the basal end of the cochlear duct and the outer compartment are more sensitive to the loss of *Fgf20* and *Sox2*, relative to the apical end and inner compartment, respectively.

**Fig 9.**
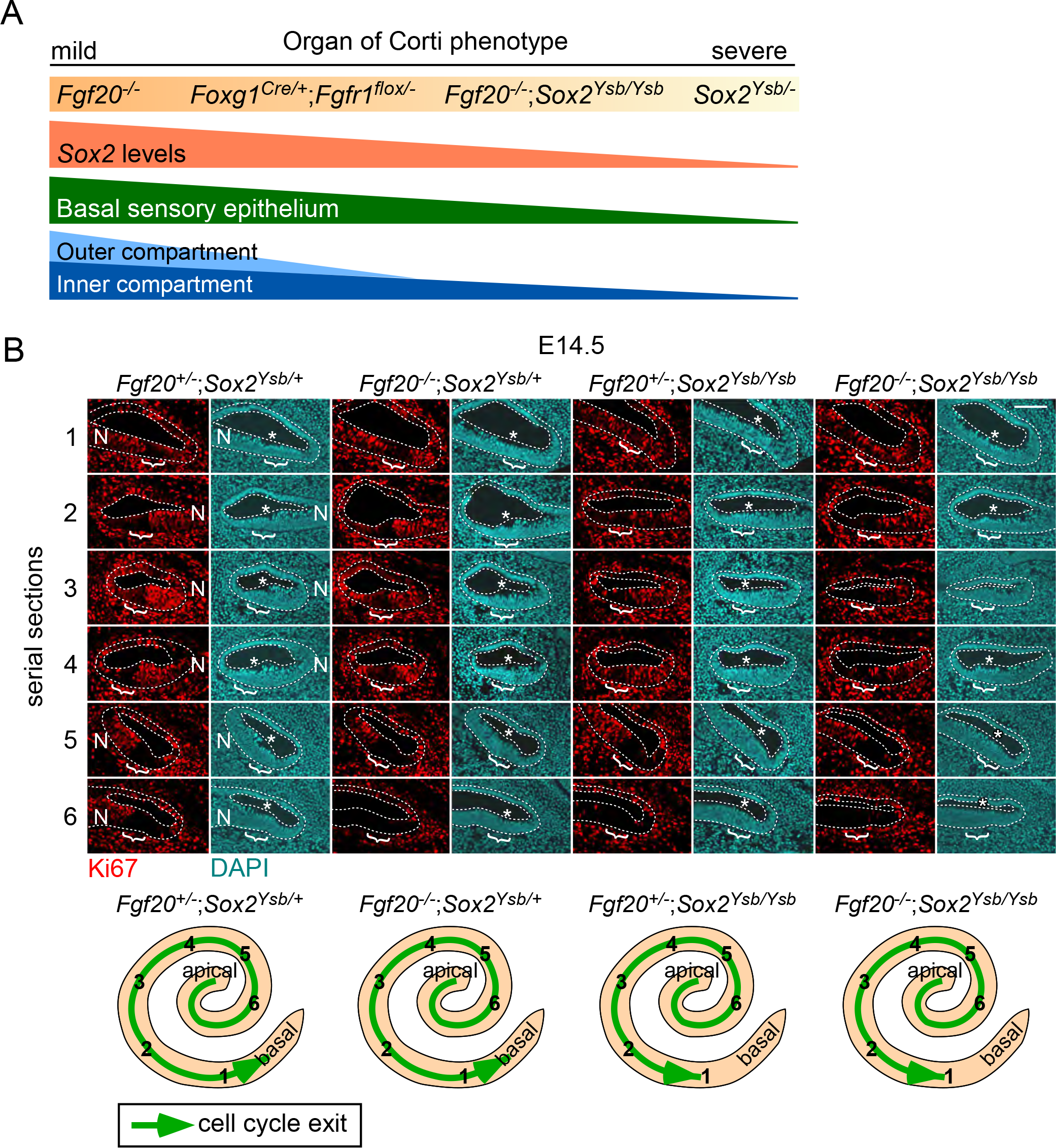
Loss of Fgf20 does not further delay prosensory specification in Sox2 hypomorph cochleae (A) The *Fgf20^-/-^*, *Foxg1^Cre/+^*;*Fgfr1^flox/-^*, *Fgf20^-/-^*;*Sox2^Ysb/Ysb^*, and *Sox2^Ysb/-^*cochleae phenotypes lie along a continuum, preferentially affecting outer and basal cochlear sensory epithelium, likely attributable to varying *Sox2* levels on an *Fgf20*-null background. (B) Serial sections (1-6) through the duct of E14.5 *Fgf20^+/-^*;*Sox2^Ysb/+^*, *Fgf20^+/-^*;*Sox2^Ysb/Ysb^*, *Fgf20^-/-^*;*Sox2^Ysb/+^*, and *Fgf20^-/-^*;*Sox2^Ysb/Ysb^*cochleae. Immunofluorescence for Ki67 (red) and DAPI (nuclei, cyan). Cochlear epithelium is outlined. Bracket indicates prosensory domain. * indicates shift of prosensory nuclei away from the luminal surface of the epithelium. N, neural side. Whole mount cochlear duct schematics show relative positions of the serial sections and progression of cell cycle exit (green arrow). Note: unlike in Fig 7, we have switched the placement of images from *Fgf20^-/-^*;*Sox2^Ysb/+^*and *Fgf20^+/-^*;*Sox2^Ysb/Ysb^* cochleae to facilitate comparison. Scale bar, 100 µm. See also S10 Fig.

To determine the mechanism underlying the *Sox2* and *Fgf20* interaction, we asked whether the similarity between the *Fgf20^-/-^*;*Sox2^Ysb/Ysb^*and *Sox2^Ysb/-^* phenotypes could be explained by a further decrease in *Sox2* levels in *Fgf20^-/-^*;*Sox2^Ysb/Ysb^*cochleae from *Sox2^Ysb/Ysb^* levels. In other words, we asked whether loss of *Fgf20* further reduces *Sox2* expression on a *Sox2* hypomorphic background. Examination of prosensory domain *Sox2* expression at E14.5 revealed, as expected, that *Fgf20^-/-^*;*Sox2^Ysb/+^*cochleae did not have a decrease in Sox2 expression compared to *Fgf20^+/-^*;*Sox2^Ysb/+^*(S10A Fig). *Fgf20^-/-^*;*Sox2^Ysb/Ysb^* cochleae also did not have a further decrease in Sox2 expression compared to *Fgf20^+/-^*;*Sox2^Ysb/Ysb^*cochleae. Moreover, despite the loss of sensory epithelium in most of the basal turn, Sox2 expression was not further decreased in the basal turn at E14.5 relative to the rest of the *Fgf20^-/-^*;*Sox2^Ysb/Ysb^*cochlea (S10A Fig). These data confirm that *Fgf20* does not regulate *Sox2* expression. A similar pattern of expression was observed for CDKN1B across the different genotypes (S10B Fig). Loss of *Fgf20* did not contribute to a further decrease in CDKN1B expression on a *Sox2^Ysb/Ysb^* background, nor was there a basal-apical difference in CDKN1B expression in *Fgf20^-^/-*;*Sox2^Ysb/Ysb^* cochleae at E14.5.

Next, we asked whether *Sox2* and *Fgf20* interact to delay prosensory specification. We showed that Fgf20-KO cochleae do not exhibit a delay in prosensory specification (Fig 6D). However, this does not rule out the possibility that the loss of *Fgf20* on a *Sox2* hypomorphic background may contribute to a delay. We examined Ki67 expression at E14.5 and found that in *Fgf20^+/-^*;*Sox2^Ysb/+^*cochleae, prosensory domain cell cycle exit has reached the end of the base (serial section 1; Fig 9B, brackets). See S10D Fig for serial “mid-modiolar” sections through the cochlea. Similarly, cell cycle exit in *Fgf20^-/-^*;*Sox2^Ysb/+^*cochleae also reached the very base. As expected, *Fgf20^+/-^*;*Sox2^Ysb/Ysb^*cochleae exhibited a slight delay in prosensory specification; cell cycle exit has reached the base (serial section 2), but has not yet reached the end of the base (serial section 1). Importantly, *Fgf20^-/-^*;*Sox2^Ysb/Ysb^* cochleae did not show a further delay relative to *Fgf20^+/-^*;*Sox2^Ysb/Ysb^*. There was also no detectable delay in basal nuclei shift in *Fgf20^-/-^*;*Sox2^Ysb/+^*or *Fgf20^-/-^*;*Sox2^Ysb/Ysb^* cochleae (Fig 9B, asterisks). These results suggest that the loss of *Fgf20* does not contribute to delayed specification and that the severity of the *Fgf20^-^/-*;*Sox2^Ysb/Ysb^* basal phenotype cannot be completely attributed to delayed specification.

Lastly, we examined proliferation in the *Fgf20^-^*and *Sox2^Ysb^* E14.5 cochleae. Interestingly, there was a noticeable decrease in the number of EdU-incorporating cells in Kölliker’s organ in *Fgf20^-^/-*;*Sox2^Ysb/Ysb^* cochleae, compared to *Fgf20^+/-^*;*Sox2^Ysb/+^*, *Fgf20^-/-^*;*Sox2^Ysb/+^*, and *Fgf20^+/-^*;*Sox2^Ysb/Ysb^* cochleae (S10C Fig). This phenotype is similar to that of *Sox2^Ysb/-^* cochleae and is less severe than that of Fgfr1-CKO cochleae. This suggests that *Fgf20* and *Sox2* interact to regulate proliferation in Kölliker’s organ, although other factors downstream of *Fgfr1* also contribute.

### Fgf20-KO organ of Corti exhibits premature differentiation

We showed that *Fgf20* likely plays a role during the initiation of differentiation. Previous studies showed that deletion of both transcription factors *Hey1* and *Hey2* results in premature differentiation in the organ of Corti [29]. Furthermore, it has been suggested that FGF signaling, in particular FGF20, regulates *Hey1* and *Hey2* expression during this process [8,29]. To test whether *Fgf20* is upstream of *Hey1* and *Hey2*, we looked at the expression of the two transcription factors via in situ hybridization. In Fgf20-KO cochleae at E14.5, *Hey1* expression is downregulated while *Hey2* is almost completely absent compared to control (Fig 10A). To test whether FGF20 loss leads to premature differentiation, we examined myosin VI (Myo6) expression, a marker of differentiated HCs [29]. At E14.5, the cochleae of 3 of 12 control embryos examined contained Myo6-expressing HCs, while the cochleae of 18 of 19 littermate Fgf20-KO embryos contained Myo6-expressing HCs (p < 0.001, Fisher’s exact test; Fig 10B). If present, the Myo6-expressing HCs at this stage were always found in the basal and mid-basal turns of the cochlea. These results show that there is premature onset of differentiation in Fgf20-KO cochleae, which begins in the basal/mid-basal turns. This result is surprising given our previous finding of delayed differentiation in the apical end of Fgf20-KO cochleae at later stages, which we confirm here (S1E Fig). These findings suggest that apical progression of differentiation may be slower in Fgf20-KO cochleae.

**Fig 10.**
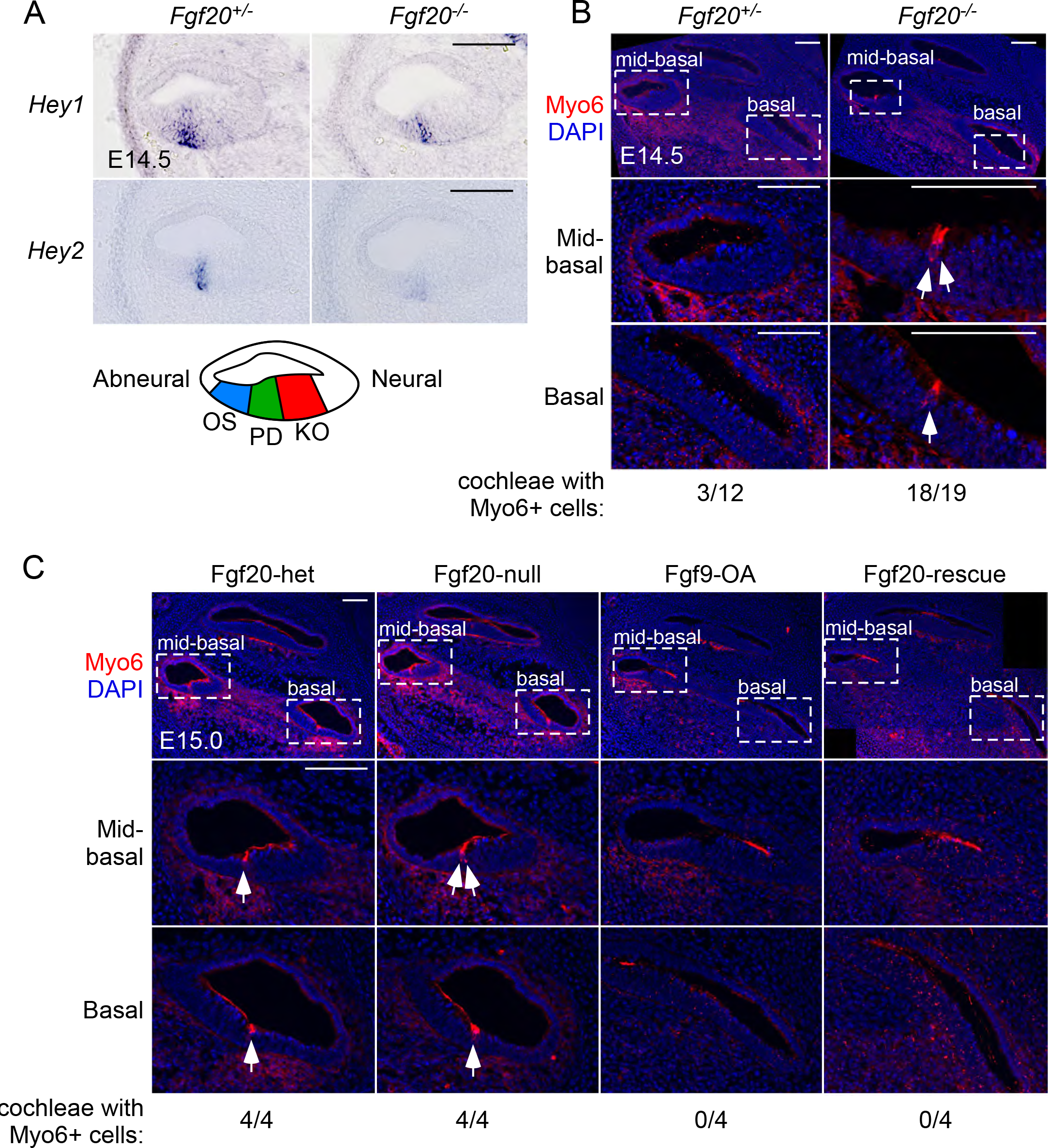
Fgf20-KO organ of Corti exhibits premature differentiation (A) RNA in situ hybridization for *Hey1* and *Hey2* on sections through the middle turn of E14.5 *Fgf20^+/-^*and *Fgf20^-/-^* cochleae. (B, C) Immunofluorescence for Myo6 (red) on “mid-modiolar” sections through the (B) E14.5 *Fgf20^+/-^* and *Fgf20^-/-^*cochleae, and (C) E15.0 Fgf20-het (*Fgf20^Cre/+^*;*ROSA^rtTA/+^*), Fgf20-null (*Fgf20^Cre/βgal^*;*ROSA^rtTA/+^*), Fgf9-OA (*Fgf20^Cre/+^*;*ROSA^rtTA/+^*;TRE-Fgf9-IRES-eGfp), Fgf20-rescue (*Fgf20^Cre/βgal^*;*ROSA^rtTA/+^*;TRE-Fgf9-IRES-eGfp) cochleae (Dox from E13.5 to E15.0), with magnification. The number of cochleae containing Myo6-expressing cells out of the total number of cochleae examined for each genotype are shown below each panel. Arrows indicate Myo6-expressing hair cells. DAPI, nuclei (blue). OS, outer sulcus; PD, prosensory domain; KO, Kölliker’s organ. Scale bar, 100 µm.

Next, we asked whether ectopic activation of FGF signaling via overexpression of FGF9 will delay the onset of differentiation. We generated Fgf20-het (*Fgf20^Cre/+^*;*ROSA^rtTA/+^*), Fgf20-null (*Fgf20^Cre/βgal^*;*ROSA^rtTA/+^*), Fgf9-OA (*Fgf20^Cre/+^*;*ROSA^rtTA/+^*;TRE-Fgf9-IRES-eGfp), Fgf20-rescue (*Fgf20^Cre/βgal^*;*ROSA^rtTA/+^*;TRE-Fgf9-IRES-eGfp) mice as before and started Dox induction at E13.5 until E15.0 (Fig 3). At E15.0, all of the Fgf20-het (4/4) and Fgf20-null (4/4) cochleae contained Myo6-expressing HCs, while none of the Fgf9-OA (0/4) and Fgf20-rescue (0/4) cochleae contained Myo6-expressing HCs (Fig 10C). This suggests that ectopic expression of FGF9 was able to delay the onset of differentiation, even with the lack of endogenous FGF20. Despite this delay in onset of differentiation, by P0, differentiation has apparently caught up in both Fgf9-OA and Fgf20-rescue cochleae (Fig 3A).

Similar to a delay in prosensory specification, premature onset of differentiation narrows the temporal buffer between the completion of specification and initiation of differentiation towards the cochlear base. In the context of a slight delay in specification due to decreased *Sox2* levels, premature differentiation from the loss of *Fgf20* can lead to an attempt at differentiation before specification in the basal end of the cochlea. We propose that *Sox2* and *Fgf20* interact to regulate the boundaries of the temporal buffer, helping to ensure that differentiation begins after the completion of specification (Fig 11).

**Fig 11.**
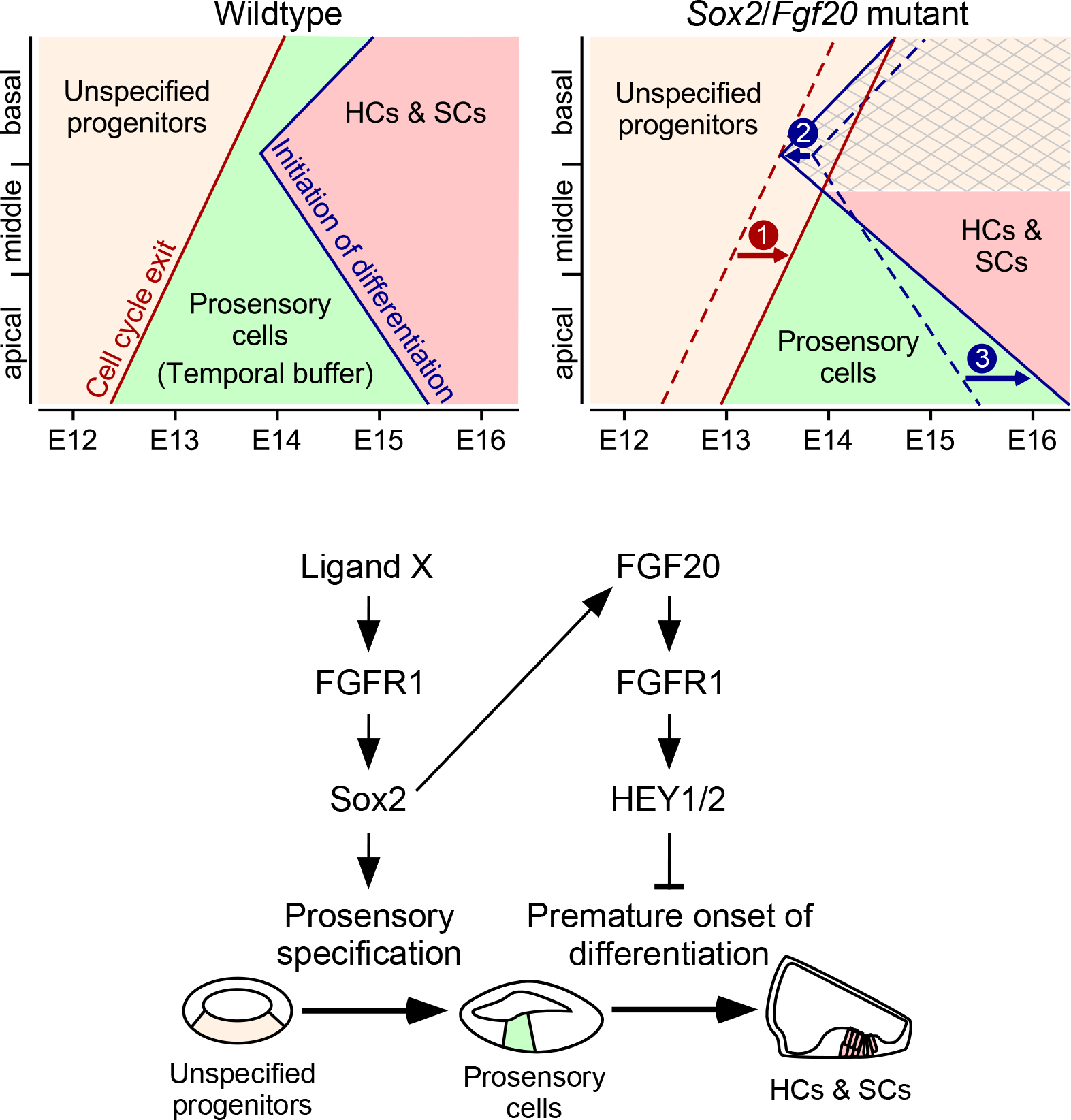
*Sox2* and *Fgf20* interact to modulate a temporal buffer between specification and differentiation Model of the roles of *Sox2* and *Fgf20* in organ of Corti development, which occurs in two stages: unspecified progenitors (tan shading) undergo specification and cell cycle exit to become prosensory cells (green shading), which then differentiate into hair cells and supporting cells (HCs & SCs; red shading). In wildtype cochleae, cell cycle exit (indicating completion of specification) begins at the apex of the cochlea and proceeds basally. Afterwards, differentiation initiates at the mid-base of the cochlea and proceeds basally and apically. The prosensory cells exist within a temporal buffer (green shading), defined as the time between cell cycle exit and initiation of differentiation. In *Sox2*/*Fgf20* mutant cochleae, decrease in levels of *Sox2* expression in the developing cochlea leads to delayed prosensory specification and cell cycle exit (arrow 1), while loss of *Fgf20* leads to premature onset of differentiation at the basal and mid-basal cochlear turns (arrow 2) as well as delayed differentiation at the apical turn (arrow 3). Loss of both *Sox2* and *Fgf20* leads to loss of the temporal buffer between specification and differentiation towards the base of the cochlear duct, disrupting the development of HCs and SCs in the basal region (crosshatch pattern).

## Discussion

### *Fgfr1* is involved in prosensory specification and differentiation, while *Fgf20* is only involved in differentiation

*Fgf20* and *Fgfr1* are required for HC and SC development. Based on similarities in the phenotype caused by the loss of FGF20 and loss of FGFR1 signaling, FGF20 has been hypothesized as the FGFR1 ligand during organ of Corti development [17–21]. However, the exact role of FGF20/FGFR1 during organ of Corti development has been a topic of debate. We previously reported that Fgf20-KO mice do not have defects in prosensory specification, and have a normally formed prosensory domain [21]. We further showed that FGF20 signaling is important during the initiation stage of differentiation, and that Fgf20-KO cochleae have gaps in the differentiated sensory epithelium filled with undifferentiated prosensory progenitors.

However, other studies have shown *in vitro* that FGF20 regulates prosensory specification via Sox2 [33] and *in vivo* that FGFR1 is required for prosensory specification via Sox2 [19]. Here, using an *in vivo* rescue model, we show that ectopic FGF9 signaling is sufficient to rescue the Fgf20-KO phenotype in a spatiotemporal pattern that matched the timing of initiation of differentiation along the length of the cochlear duct. We conclude, therefore, that FGF20 is involved in differentiation and not necessary for prosensory specification.

Notably, the Fgf20-KO phenotype, in which two-thirds of OHCs fail to develop, is not as severe as the Fgfr1-CKO phenotype, which lacks almost all OHCs as well as half of IHCs. Potential explanations for this include differences in mouse genetic background, and the existence of a redundant FGF ligand(s). To rule out the former, we examine here Fgf20-KO and Fgfr1-CKO mice on a similar genetic background, and replicated the difference in phenotype severity. We also replicated the decrease in *Sox2* expression in the prosensory domain previously reported in Fgfr1-CKO mice [19]. We further reaffirmed that *Sox2* expression in the prosensory domain is not affected by the loss of *Fgf20*. This suggests that another FGF ligand signaling through FGFR1 is required to maintain *Sox2* expression during prosensory specification. The identity of this ligand is currently unknown.

*Foxg1^Cre^* has been used in several studies to target the otic epithelium, including to conditionally delete *Fgfr1* [18–20]. One concern with *Foxg1^Cre^* is that it is a null allele [22]. *Foxg1*-null mice have shortened cochlear length, although HC and SC differentiation did not appear to be directly affected [34]. Previous work [35] and our results here showed that *Foxg1* is not haploinsufficient during cochlea development, as *Foxg1^Cre/+^*;*Fgfr1^flox/+^*cochleae had very similar phenotypes to *Fgfr1^+/-^* cochleae. Moreover, the use of the Six1enh21-Cre transgene, which targets the otic epithelium in a similar spatiotemporal pattern as *Foxg1^Cre^*, to conditionally delete *Fgfr1* resulted in the same phenotype as *Foxg1^Cre/+^*;*Fgfr1^flox/-^*cochleae [19]. This included the loss of almost all OHCs, loss of IHCs, and decreased prosensory *Sox2* expression. Therefore, the increased severity of *Foxg1^Cre/+^*;*Fgfr1^flox/-^*cochleae relative to *Fgf20^-/-^* cochleae is likely not attributable *Foxg1* haploinsufficiency.

We hypothesized that the severity of the Fgfr1-CKO phenotype is due to the loss of FGF20 signaling during differentiation and decreased *Sox2* expression causing disrupted prosensory specification. Consistent with this hypothesis, the combination of *Fgf20^-/-^*and *Sox2^Ysb/Ysb^* mutations phenocopied Fgfr1-CKO cochleae. The similarities in phenotype include approximately a 30% reduction in cochlear length and almost a complete loss of OHCs and SCs and approximately a 50% loss of IHCs. Interestingly, the *Fgf20^-/-^*;*Sox2^Ysb/Ysb^*phenotype is also similar to the *Sox2^Ysb/-^* phenotype. We conclude that the Fgfr1-CKO, *Fgf20^-/-^*;*Sox2^Ysb/Ysb^*, and *Sox2^Ysb/-^* phenotypes likely lie along the same continuum, as these three genotypes all exhibited a lack of *Fgf20* expression or signaling and varying levels of *Sox2* expression (Fig 9A). Fgf20-KO cochleae, in which *Sox2* expression was not affected, lies at the mild end of this continuum. Interestingly, this continuum shows that in the absence of *Fgf20* expression or signaling, reductions in the level of *Sox2* most severely affected sensory epithelium development of the cochlear base and the outer compartment. Moving from the Fgf20-KO (mild) end of the spectrum towards the *Sox2^Ysb/-^*(severe) end, increasing numbers of HCs and SCs are lost, preferentially form the cochlear base and the outer compartment.

### *Sox2* and *Fgf20* interact to affect development towards the basal end of the cochlea

We show here conclusive evidence that *Sox2* and *Fgf20* genetically interact during cochlea development. Interestingly, HC and SC development towards the basal end of the cochlea is more severely affected by the loss of *Sox2* and *Fgf20* and their interaction. While we hypothesize that *Sox2* and *Fgf20* are involved in distinct steps during organ of Corti development (prosensory specification and differentiation, respectively), there is nevertheless potential for a strong interaction. We propose that the timing of specification and differentiation define a temporal buffer that normally prevents differentiation from initiating prior to the completion of specification, and that *Sox2* and *Fgf20* modulate the borders of this buffer. In a developmental pathway, the upstream event (specification) must occur prior to the downstream event (differentiation). Therefore, loss of *Sox2* and *Fgf20* leading to delayed specification and premature differentiation onset, respectively, disrupts the temporal buffer, especially towards the cochlear base (Fig 11).

Specification must occur prior to the onset of differentiation. However, cell cycle exit does not need to occur prior to the onset of differentiation, as mice lacking *Cdkn1b*, which is required for cell cycle exit, still produce HCs and SCs (Chen & Segil, 1999; Kanzaki et al., 2006). Here, we use cell cycle exit in the prosensory domain (also known as the zone of non-proliferation) as a marker for the completion of specification (Chen & Segil, 1999). We hypothesize that prosensory cells become specified and primed for differentiation upon withdrawal from the cell cycle. Previous studies showed that prosensory cells are indeed capable of differentiating into HCs and SCs directly after cell cycle exit, even in the apex. When *Shh* was deleted from the spiral ganglion, differentiation began in the apex shortly after cell cycle exit and progressed towards the base [5]. This suggests that specification occurs in an apex-to-base direction. We cannot rule out, however, that specification occurs in the same direction as differentiation (base-to-apex), independently of cell cycle exit. Such a scenario would still be consistent with our model that a combination of delayed specification and premature onset of differentiation accounts for the more severe basal phenotype in *Fgf20/Sox2* mutants.

The effect of loss of *Fgf20* on the timing of differentiation is small. We estimate that the onset of differentiation in Fgf20-KO cochleae is advanced by only around 0.5 days. By itself, this effect does not lead to a more severe mid-basal or basal phenotype in Fgf20-KO cochleae. However, we present evidence that on a sensitized genetic background of delayed specification, this small change in the timing of differentiation leads to a large defect in HC and SC production towards the basal end of the cochleae. We propose that this at least partially explains the interaction between *Sox2* and *Fgf20*. Furthermore, the relative sparing of development towards the apical end of *Sox2^Ysb/Ysb^*;*Fgf20^-/-^*cochleae, especially of IHCs, can be further explained by a delay in differentiation at the apical end due to the loss of *Fgf20*. We do not know why an apical-basal difference in timing of differentiation exists in Fgf20-KO cochleae. Perhaps there is a delay in the apical progression of differentiation, or perhaps other factors contribute to the differentiation of the apical end of the cochlea. Consistent with the latter, by P7 in Fgf20-KO cochleae, the apical tip contains a full complement of IHCs and OHCs [21].

Notably, while we show the potential for a *Sox2* and *Fgf20* interaction in modulating the temporal buffer between specification and differentiation, *Sox2* also has known roles during HC and SC differentiation [13,15,37,38]. Therefore, the genetic interaction may occur during differentiation as well. While interaction at this stage may explain the preferential loss of outer compartment cells in *Sox2* and *Fgf20* mutants, it does not explain the selective loss of basal cochlear HCs and SCs. Therefore, we conclude that the *Sox2* and *Fgf20* interaction regulates the temporal buffer, with potential further interactions during differentiation.

The Notch ligand Jagged1 (Jag1) is thought to be important for cochlear prosensory specification via lateral induction [39–45]. Interestingly, Notch signaling has also been shown to be upstream of both *Fgf20* and *Sox2* in the developing cochlea [33]. Conditional deletion of *Jag1* or *Rbpj*, the major transcriptional effector of canonical Notch signaling, resulted in the loss of HCs and SCs, particularly from the basal end of the cochlear duct, similarly to *Fgf20*/*Sox2* mutants. Unlike *Fgf20*/*Sox2* mutants, however, deletion of *Jag1* or *Rbpj* led to preferential loss of Sox2 and CDKN1B expression from the prosensory domain at the basal end of the cochlea [40,42,46]. This suggests that Jag1-Notch signaling is required for prosensory specification, especially towards the cochlear base. This likely accounts for the more severe basal phenotype of *Jag1* or *Rbpj* mutants. This same mechanism likely does not explain the more severe basal phenotype of *Fgf20/Sox2* mutants, as Sox2 and CDKN1B expression was not more severely reduced or absent in the cochlear base in these mice. Notably, not all studies agree that *Jag1* or *Rbpj* is required for Sox2 and CDKN1B expression or for prosensory specification [47]. Further studies are required to elucidate the functional relationship between *Jag1/Notch*, *Fgf20*, and *Sox2* during cochlea development.

Other genes that potentially interact with *Fgf20* and *Sox2* during cochlea development include *Mycn* (*N-Myc*) and *Mycl* (*L-Myc*). Interestingly, deletion of *Mycn* and *Mycl* from the cochlear epithelium results in accelerated cell cycle exit and delayed initiation of differentiation [48], opposite to the effects of loss of *Sox2* and *Fgf20*. Addressing potential interactions between *Sox2*, *Fgf20*, *Mycn*, and *Mycl* is another topic for future studies.

### Outer compartment of the cochlear sensory epithelium is more sensitive to the loss of *Fgfr1, Fgf20*, and *Sox2* than the inner compartment

In all of the genotypes we observed in this study, the loss of outer compartment cells (i.e. OHCs) was predominant. Only in the most severe cases in which almost all OHCs were missing, as seen in Fgfr1-CKO, *Fgf20^-/-^*;*Sox2^Ysb/Ysb^*, and *Sox2^Ysb/-^* cochleae, were IHCs also lost. Similarly, reduction in SC number always preferentially affected the outermost cells. This suggests that the organ of Corti outer compartment is more sensitive to the loss of *Fgfr1*, *Fgf20,* and *Sox2* than the inner compartment. The combination of *Fgf20^-^* and *Sox2^Ysb^* alleles elegantly demonstrates this: as the number of *Fgf20^-^* and *Sox2^Ysb^*alleles increased, the number of OHCs progressively decreased. In the double homozygous mutants, the number of IHCs decreased as well.

Previous studies noted that the dosage of *Fgfr1* affects the degree of organ of Corti outer compartment loss. In *Fgfr1* hypomorphs with 80% reduction in transcription, only the third row of OHCs were missing, while 90% hypomorphs had a slightly more severe phenotype [20]. Therefore, *Fgfr1* loss preferentially affects the outermost HCs. Other studies suggested that the timing of *Fgfr1* deletion is important in determining the degree of outer compartment loss and level of *Sox2* expression. When an earlier-expressed Cre driver (Six1enh21-Cre) was used to conditionally delete *Fgfr1*, almost all OHCs and some IHCs were lost, with a 66% reduction in *Sox2* expression at E14.5 [19]. When a later-expressed Cre driver (*Emx2^Cre^*) was used, many more OHCs and IHCs remained, with only a 12% reduction in *Sox2* expression. Our results are consistent with both of these studies. We show that FGF20-independent FGFR1 signaling and *Sox2* are required early, affecting both IHC and OHC development, while FGF20-FGFR1 signaling is important during later stages, affecting only OHC development.

Differentiation in the organ of Corti not only occurs in a basal-to-apical gradient, but also occurs in an orthogonal inner-to-outer gradient. That is, IHCs differentiate first, followed by each sequential row of OHCs [49]. This wave of differentiation suggests that perhaps outer compartment HCs and SCs require a longer temporal buffer between specification and differentiation. The genetic interaction between *Sox2* and *Fgf20* in modulating this temporal buffer, therefore, could also account for the loss of outer compartment HCs and SCs. We hypothesize that the requirement for a longer temporal buffer may also be involved in determining OHC fate. In *Fgf20^+/-^*;*Sox2^+/-^*cochleae, there was a slight decrease in OHCs that was compensated for by ectopic IHCs, suggesting a fate switch from OHCs into IHCs. Here, we confirmed previous suggestions that *Fgf20* regulates *Hey1* and *Hey2* to prevent premature differentiation in the developing organ of Corti [8,29]. Interestingly, in *Hey1*/*Hey2* double knockout cochleae, there was a similar slight decrease in OHCs compensated for by ectopic IHCs [29]. Furthermore, inner ear-specific deletion of either *Smoothened* or *Neurod1*, which led to premature differentiation in the apical cochlear turn, also led to loss of OHCs and the presence of ectopic IHCs at the apex [6,8]. These findings further support a model where timing of specification and differentiation affect IHC versus OHC fate, an interesting and important topic for future studies.

Previously, we hypothesized that *Fgf20* is strictly required for the differentiation of an outer compartment progenitor [21]. However, data we present here show that *Fgf20*, on a sensitized, *Sox2* hypomorphic background, is also required for inner compartment differentiation. We conclude that inner and outer compartment progenitors likely are not distinct populations. Rather, all prosensory progenitors giving rise to the organ of Corti exist on an inner-to-outer continuum. FGF20 signaling, in combination with other factors including Sox2, are required for the proper development of all of these cells, though with varying sensitivities.

### The relationship between *Fgf20* and *Hey1/Hey2* in regulating differentiation is complex

We show *in vivo* that *Fgf20* is upstream of *Hey1* and *Hey2*. Supporting this result, *Fgfr1* has also been shown *in vivo* to be upstream of *Hey2* [19]. Interestingly, in explant studies, inhibition of FGF signaling alone did not result in decreased *Hey1/Hey2* expression or premature differentiation [29]. However, FGF inhibition was able to rescue the overexpression of *Hey1*/*Hey2* and the delay in differentiation induced by SHH signaling overactivation [8,29]. These discrepancies suggest that the relationship between *Fgf20* and *Hey1*/*Hey2* is more complicated than we currently understand. Notably, *Hey1*/*Hey2* double knockout cochleae do not exhibit a loss of OHCs to the extent of Fgf20-KO cochleae, suggesting that other genes downstream of *Fgf20* are important in organ of Corti development. Moreover, deletion of *Fgf20* only led to premature differentiation at the basal and mid-basal turns. *Fgf20* deletion actually delayed differentiation in the apical end of the cochlea. Deletion of *Hey1*/*Hey2*, contrarily, led to premature differentiation along the entire length of the cochlear duct, although it is unclear how *Hey1/Hey2* loss affects the timing of apical differentiation beyond E15.0 [29]. This suggests that other factors downstream of *Fgf20* interact with *Hey1*/*Hey2* to regulate the timing of differentiation. Perhaps these same genes contribute to the loss of OHCs in Fgf20-KO cochleae. *Mekk4*, which has been shown to be downstream of *Fgf20* and necessary for OHC differentiation [50] could be one of these genes. Identifying other factors downstream of *Fgf20* will be a topic of future studies.

## Materials and methods

**Key Resources Table.**

| Reagent type<br>(species) or<br>resource | Designation | Source or<br>reference | Identifiers | Additional<br>informatio<br>n |
| --- | --- | --- | --- | --- |
| Genetic reagent<br>(*M. musculus*) | *Fgf20*<sup>Cre</sup> | Huh et al., 2015 | MGI:5751785 |  |
| Genetic reagent<br>(*M. musculus*) | *Fgf20*<sup>βgal</sup> | Huh et al., 2012 | RRID:MGI:5425887 |  |
| Genetic reagent<br>(*M. musculus*) | *Foxg1*<sup>Cre</sup> | Hébert and<br>McConnell,<br>2000 | RRID:IMSR_JAX:00<br>4337 |  |
| Genetic reagent<br>(*M. musculus*) | *Fgfr1*<sup>flox</sup> | Trokovic et al.,<br>2003 | RRID:IMSR_CMMR:<br>0268 |  |
| Genetic reagent<br>(*M. musculus*) | *Sox2*<sup>Cre</sup> | Hayashi et al.,<br>2002 | RRID:IMSR_JAX:00<br>4783 |  |
| Genetic reagent<br>(*M. musculus*) | *Sox2*<sup>Ysb</sup> | Dong et al.,<br>2002 | RRID:IMSR_EM:050<br>15 |  |
| Genetic reagent<br>(*M. musculus*) | *Sox2*<sup>flox</sup> | Shaham et al.,<br>2009 | RRID:IMSR_JAX:01<br>3093 |  |
| Genetic reagent<br>(*M. musculus*) | *ROSA*<sup>rtTA</sup> | Belteki et al.,<br>2005 | RRID:IMSR_JAX:00<br>5670 |  |
| Genetic reagent<br>(*M. musculus*) | TRE-Fgf9-IRES-eGfp | White et al.,<br>2006 | MGI:5538516 |  |
| Recombinant<br>DNA reagent | *pBluescriptKS-Fgfr1TM* | K. Peters |  |  |
| Recombinant<br>DNA reagent | *pGEMT-Fgf20* |  |  |  |
| Recombinant<br>DNA reagent | *pBluescriptSK-Sox2* | R. Lovell-Badge |  | Gift of A.<br>Kiernan |
| Recombinant<br>DNA reagent | *pGEM-Etv4* | G. Martin |  |  |
| Recombinant<br>DNA reagent | *pBluescriptSK-Etv5* | G. Martin |  |  |
| Recombinant<br>DNA reagent | *pT7T3D-Hey1* | IMAGE clone | #478014 | Gift of S.<br>Rentschler |
| Recombinant<br>DNA reagent | *pCMVSPORT6-Hey2* | IMAGE clone | #5374813 | Gift of S.<br>Rentschler |
| Antibody | Sheep polyclonal<br>anti-Digoxigenin-AP | Sigma-Aldrich | 11093274910 | 1:1000 |
| Antibody | Rabbit polyclonal<br>anti-P75NTR | EMD Millipore | AB1554 | 1:300 |
| Antibody | Rabbit polyclonal<br>anti-Prox1 | EMD Millipore | ABN278 | 1:1000 |
| Antibody | Goat polyclonal<br>anti-Sox2 | Santa Cruz<br>Biotechnology | sc-17320 | 1:200 |
| Antibody | Rabbit polyclonal anti-p27Kip1 | Neomarkers | RB-9019-P | 1:50 |
| Antibody | Rabbit polyclonal anti-Ki67 | Abcam | ab15580 | 1:200 |
| Antibody | Rabbit polyclonal anti-Myo6 | Santa Cruz Biotechnology | sc-50461 | 1:100 |
| Antibody | Donkey polyclonal anti-Rabbit IgG, Alexa Fluor 488 | Thermo-Fisher Scientific | A-21206 | 1:500 |
| Antibody | Donkey polyclonal anti-Goat IgG, Alexa Fluor 555 | Thermo-Fisher Scientific | A-21432 | 1:500 |
| Antibody | Goat polyclonal anti-Rabbit IgG, Flexa Fluor 555 | Thermo-Fisher Scientific | A-21428 | 1:500 |
| Chemical compound, drug | Alexa Fluor 488-conjugated Phalloidin | Invitrogen | A12379 | 1:50 |
| Chemical compound, drug | Dox Diet, Grain-Based Doxycycline | Bio-Serv | S3888 |  |
| Chemical compound, drug | EdU (5-ethynyl-2'-deoxyuridine) | Thermo-Fisher Scientific | E10187 |  |
| Commercial assay or kit | Click-iT EdU Alexa Fluor 488 kit | Thermo-Fisher Scientific | C10337 |  |

### Mice

Mice were group housed with littermates, in breeding pairs, or in a breeding harem (2 females to 1 male), with food and water provided ad libitum.

For timed-pregnancy experiments, embryonic day 0.5 (E0.5) was assigned as noon of the day the vaginal plug was found. For postnatal experiments, postnatal day 0 (P0) was determined as the day of birth.

Mice were of mixed sexes and maintained on a mixed C57BL/6J × 129X1/SvJ genetic background. All mice were backcrossed at least three generations onto this background. The following mouse lines were used:

- *Fgf20^Cre^* (*Fgf20^-^*): knockin allele containing a sequence encoding a GFP-Cre fusion protein replacing exon 1 of *Fgf20*, resulting in a null mutation [18].
- *Fgf20^βgal^*: knockin allele containing a sequence encoding β-galactosidase (βgal) replacing exon 1 of *Fgf20*, resulting in a null mutation [21].
- *Foxg1^Cre^*: knockin allele containing a sequence encoding Cre fused in-frame downstream of the first 13 codons, resulting in a null mutation [22].
- *Fgfr1^flox^*: allele containing loxP sequences flanking exons 8 through 15 of *Fgfr1*. Upon Cre-mediated recombination, produces a null mutation [51]
- *Fgfr1^-^*: null allele generated by combining *Fgfr1*^flox^ with Sox2^Cre^ [52] to delete *Fgfr1* from the epiblast.
- *ROSA^rtTA^*: knockin allele containing a loxP-Stop-loxP sequence followed by a sequence encoding rtTA-IRES-eGFP, targeted to the ubiquitously expressed *ROSA26* locus. Upon Cre-mediated recombination, reverse tetracycline transactivator (rtTA) and eGFP are expressed [25].
- TRE-Fgf9-IRES-eGfp: transgene containing seven tetracycline-inducible regulatory elements driving the expression of FGF9-IRES-eGFP [26].
- *Sox2^Ysb^*: Inner ear specific *Sox2* hypomorphic allele resulting from a random insertion of a transgene in chromosome 3, likely interfering with tissue-specific *Sox2* regulatory elements [11].
- *Sox2^-^*: null allele generated by combining *Sox2^flox^* [53] with Sox2^Cre^ to delete *Sox2* from the epiblast.

All studies performed were in accordance with the Institutional Animal Care and Use Committee at Washington University in St. Louis (protocol #20160113) and University of Nebraska Medical Center (protocol #16-004-02 and 16-005-02).

### Doxycycline induction

Pregnant dams were starved overnight the night before initiation of Dox induction and fed Dox Diet, Grain-Based Doxycycline, 200 mg/kg (S3888, Bio-Serv, Flemington, NJ) ad libitum starting at noon on the start date of Dox induction. On the stop date of Dox induction, Dox Diet was replaced with regular mouse chow at noon.

### Sample preparation and sectioning

For whole mount cochleae, inner ears were dissected out of P0 pups and fixed in 4% PFA in PBS overnight at 4°C with gentle agitation. Samples were then washed x3 in PBS. Cochleae were dissected away from the vestibule, otic capsule, and periotic mesenchyme with Dumont #55 Forceps (RS-5010, Roboz, Gaithersburg, MD). The roof of the cochlear duct was opened up by dissecting away the stria vascularis and Reissner’s membrane; tectorial membrane was removed to expose hair and supporting cells.

For sectioning, heads from E14.5 embryos were fixed in 4% PFA in PBS overnight at 4°C with gentle agitation. Samples were then washed x3 in PBS and cryoprotected in 15% sucrose in PBS overnight and then in 30% sucrose in PBS overnight. Samples were embedded in Tissue-Tek O.C.T. compound (4583, VWR International, Radnor, PA) and frozen on dry ice. Serial horizontal sections through base of the head were cut at 12 µm with a cryostat, dried at room temperature, and stored at −80°C until use.

### RNA in situ hybridization

Probe preparation: mouse cDNA plasmids containing the following inserts were used to make RNA in situ probes, and were cut and transcribed with the indicated restriction enzyme (New England Biolabs, Ipswich, MA) and RNA polymerase (New England Biolabs, Ipswich, MA): *Fgfr1* transmembrane domain (325 bp, HincII, T7, gift of K. Peters), *Fgf20* (653 bp, NcoI, Sp6), *Sox2* (750 bp, AccI, T3, gift of A. Kiernan), *Etv4* (∼2300 bp, ApaI, Sp6, gift of G. Martin), *Etv5* (∼4000 bp, HindIII, T3, gift of G. Martin), *Hey1* (343 bp, EcoRI, T3, gift of S. Rentschler), *Hey2* (819 bp, EcoRI, T7, gift of S. Rentschler). Restriction digest and *in vitro* transcription were done according to manufacturer’s instructions, with DIG RNA Labeling Mix (11277073910, Sigma-Aldrich, St. Louis, MO). After treatment with RNase-free DNase I (04716728001, Sigma-Aldrich, St. Louis, MO) for 15 min at 37°C, probes were hydrolyzed in hydrolysis buffer (40 mM NaHCO_3_, 60 mM Na_2_CO_3_) at 60°C for up to 30 min, depending on probe size.

Frozen section in situ hybridization: frozen slides were warmed for 20 min at room temperature and then 5 min at 50°C on a slide warmer. Sections were fixed in 4% PFA in PBS for 20 min at room temperature, washed x2 in PBS and treated with pre-warmed 10 µg/ml Proteinase K (03115828001, Sigma-Aldrich, St. Louis, MO) in PBS for 7 min at 37°C. Sections were then fixed in 4% PFA in PBS for 15 min at room temperature, washed x2 in PBS, acetylated in 0.25% acetic anhydrate in 0.1M Triethanolamine, pH 8.0, for 10 min, and washed again in PBS. Sections were then placed in pre-warmed hybridization buffer (50% formamide, 5x SSC buffer, 5 mM EDTA, 50 µg/ml yeast tRNA) for 3 h at 60°C in humidified chamber for prehybridization. Sections were then hybridized in 10 µg/ml probe/hybridization buffer overnight (12-16 h) at 60°C. The next day, sections were washed in 1x SSC for 10 min at 60°C, followed by 1.5x SSC for 10 min at 60°C, 2x SSC for 20 min at 37°C x2, and 0.2x SSC for 30 min at 60°C x2. Sections were then washed in KTBT (0.1 M Tris, pH 7.5, 0.15 M NaCl, 5 mM KCl, 0.1% Triton X-100) at room temperature and blocked in KTBT + 20% sheep serum + 2% Blocking Reagent (11096176001, Sigma-Aldrich, St. Louis, MO) for 4 h. Blocking Reagent was dissolved in 100 mM Maleic acid, 150 mM NaCl, pH 7.5. Sections were then incubated in sheep anti-Digoxigenin-AP, Fab fragments (1:1000, 11093274910, Sigma-Aldrich, St. Louis, MO) in KTBT + 20% sheep serum + 2% Blocking Reagent overnight at 4°C. Sections were then washed x3 in KTBT for 30 min at room temperature, and then washed x2 in NTMT (0.1 M Tris, pH 9.5, 0.1 M NaCl, 50 mM MgCl_2_, 0.1% Tween 20) for 15 min. Sections were next incubated in NTMT + 1:200 NBT/BCIP Stock Solution (11681451001, Sigma-Aldrich, St. Louis, MO) in the dark at room temperature until color appeared. Sections were then washed in PBS, post-fixed in 4% PFA in PBS for 15 min and washed x2 in PBS. Finally, sections were dehydrated in 30% and then 70% methanol, 5 min each, followed by 100% methanol for 15 min. Sections were then rehydrated in 70% and 30% methanol and then PBS, 5 min each, and mounted in 95% glycerol.

### Immunofluorescence

Whole mount: cochleae were incubated in PBS + 0.5% Tween-20 (PBSTw) for 1 h to permeabilize. Cochleae were then blocked using PBSTw + 5% donkey serum for 1 h and then incubated in PBSTw + 1% donkey serum with the primary antibody overnight at 4°C. Cochleae were then washed x3 in PBS and incubated in PBS + 1% Tween-20 with the secondary antibody. After wash in PBS x3, cochleae were mounted in 95% glycerol with the sensory epithelium facing up.

Frozen slides were warmed for 30 min at room temperature and washed in PBS before incubating in PBS + 0.5% Triton X-100 (PBST) for 1 h to permeabilize the tissue. Sections were then blocked using in PBST + 5% donkey serum for 1 h and then incubated in PBST + 1% donkey serum with the primary antibody overnight at 4°C in a humidified chamber. Sections were then washed x3 in PBS and incubated in PBS + 1% Triton X-100 with the secondary antibody. After wash in PBS x3, slides were mounted in VectaShield antifade mounting medium with DAPI (H-1200, Vector Labs, Burlingame, CA).

### Cell proliferation assay

EdU (E10187, Thermo-Fisher Scientific, Waltham, MA) was injected i.p. into pregnant dams at 100 µg per gram body weight. Embryos were harvested at 1 h after injection. EdU was detected using the Click-iT EdU Alexa Fluor 488 kit (C10337, Thermo-Fisher Scientific, Waltham, MA) according to manufacturer’s instructions.

### Imaging

Brightfield microscopy was done using a Hamamatsu NanoZoomer slide scanning system with a 20x objective. Images were processed with the NanoZoomer Digital Pathology (NDP.view2) software.

Fluorescent microscopy was done using a Zeiss LSM 700 confocal or Zeiss Axio Imager Z1 with Apotome 2, with z-stack step-size determined based on objective lens type (10x or 20x), as recommended by the ZEN software (around 1 µm). Fluorescent images shown are maximum projections. Low magnification fluorescent images shown of the whole cochlear duct required stitching together, by hand, several images. Images were processed with ImageJ (imagej.nih.gov).

### Quantification

Measurements and cell quantification (using the Cell Counter plugin by Kurt De Vos) were done using ImageJ. Total cochlear duct length was defined as the length from the very base of the cochlea to the very tip of the apex, along the tunnel of Corti. Hair cells were identified via Phalloidin, which binds to F-actin [54]. Inner pillar cells were labeled via p75NTR [55], and supporting cells (SCs, including pillar cells and Deiters’ cells) were labeled with a combination of Prox1 [56] and Sox2 [10]. Inner hair cells (IHCs) were differentiated from outer hair cells (OHCs) based on their neural/abneural location, respectively, relative to p75NTR-expressing inner pillar cells. For total cell counts, IHCs, OHCs, and SCs were counted along the entire length of the cochlea. Total cell counts were also normalized to cochlear length and presented as cell count per 100 µm of cochlea (e.g. IHCs/100 µm). For cell quantification at the basal, middle, and apical turns of the cochlea, the cochlear duct was evenly divided into thirds, and total IHCs, OHCs, and SCs were quantified for each third and normalized to length. For the Fgf20-rescue experiments in Fig 3, IHCs, OHCs, and SCs from at least 300 µm regions of the basal (10%), middle (40%), and apical (70%) turns of the cochleae were counted and normalized to 100 µm along the length of the cochlear duct.

In Sox2^Ysb/-^ cochleae, p75NTR expression was mostly absent, resulting in sensory islands without p75NTR-expressing inner pillar cells. In these cochleae, HCs not associated with inner pillar cells were presumed to be IHCs during quantification. When a curved line was drawn connecting the p75NTR islands along the organ of Corti, these presumed IHCs were always medial (neural) to that line.

### Statistical analysis and plotting

All figures were made in Canvas X (ACD systems). Data analysis was performed using the Python programming language (python.org) in Jupyter Notebook (jupyter.org) with the following libraries: Pandas (pandas.pydata.org), NumPy (numpy.org) and SciPy (scipy.org). Plotting was done using the Matplotlib library (matplotlib.org). Statistics (t-test, one-way ANOVA, two-way ANOVA, and Fisher’s exact test) were performed using the SciPy module Stats; Tukey’s HSD was performed using the Statsmodels package (statsmodels.org). All comparisons of two means were performed using two-tailed, unpaired Student’s t-test. For comparisons of more than two means, one-way ANOVA was used, except in Fig 8 and S7 Fig, where two-way ANOVA was used, with the factors being *Fgf20* (levels: *Fgf20^+/+^*, *Fgf20^+/-^*, *Fgf20^-/-^*) and *Sox2* (levels: *Sox2^+/+^*, *Sox2^Ysb/+^*, *Sox2^Ysb/Ysb^*) gene dosage. For significant ANOVA results at α = 0.05, Tukey’s HSD was performed for post-hoc pair-wise analysis. In all cases, p < 0.05 was considered statistically significant. All statistical details can be found in the figures and figure legends. In all cases, each sample (each data point in graphs) represents one animal. Based on similar previous studies, a sample size of 3-5 was determined to be appropriate. Error bars represent mean ± standard deviation (SD). For qualitative comparisons (comparing expression via immunofluorescence or RNA in situ hybridization), at least three samples were examined per genotype. All images shown are representative.

Evaluation of onset of Myo6-expressing cells (Figs 10B and 10C): 3 or 4 serial sections through the entire cochleae were immunostained for Myo6 and evaluated, blinded to genotype, for the presence of Myo6-expressing cells. E14.5 embryos were further stage-matched based on interdigital webbing of the hindlimb (at E14.5, roughly half of the hindlimb interdigital webbing is still present). Of the 34 embryos at E14.5, 3 were removed from analysis due to lack of or minimal hindlimb interdigital webbing (too old relative to the other embryos).

## Acknowledgements

We would like to thank Drew Hagan, Kel Vin Woo, and Yongjun Yin for their critical reading of the manuscript.

## Supplemental figure titles and legends

**S1 Fig.**

(A-C) Quantification of length-normalized number of (A) inner hair cells (IHCs/100 µm), (B) outer hair cells (OHCs/100 µm), and (C) supporting cells (SCs/100 µm) in the basal, middle, and apical turns of P0 cochleae from *Fgf20^+/-^*, *Fgf20^-/-^ Fgfr1^flox/+^*, *Fgfr1^flox/-^*, *Foxg1^Cre/+^*;*Fgfr1^flox/+^*, and *Foxg1^Cre/+^*;*Fgfr1^flox/-^* mice. *Fgf20^+/-^*and *Fgf20^-/-^* cochleae were analyzed by unpaired Student’s t test; *Fgfr1^flox/+^*, *Fgfr1^flox/-^*, *Foxg1^Cre/+^*;*Fgfr1^flox/+^*, and *Foxg1^Cre/+^*;*Fgfr1^flox/-^* cochleae were analyzed by one-way ANOVA. P values shown are from the t test and ANOVA. * indicates p < 0.05 from Student’s t test or Tukey’s HSD (ANOVA post-hoc); n.s., not significant. Error bars, mean ± SD.

(D) Schematic showing the positions of basal, middle, and apical turns along the cochlear duct. Apical tip refers to the apical end of the cochlea.

(E) Whole mount cochlea from P0 *Fgf20^+/-^*, *Fgf20^-/-^*, *Foxg1^Cre/+^*;*Fgfr1^flox/+^*, and *Foxg1^Cre/+^*;*Fgfr1^flox/-^* mice showing immunofluorescence for phalloidin (green) and p75NTR (red) at the apical tip of the cochlea. T, towards the tip. Scale bar, 100 µm.

**S2 Fig.**

(A, B) Sections through the middle turn of E14.5 cochlear ducts from *Fgf20^+/-^*, *Fgf20^-/-^*, *Foxg1^Cre/+^*;*Fgfr1^flox/+^*, and *Foxg1^Cre/+^*;*Fgfr1^flox/-^* mice. Scale bar, 100 µm. Refer to schematic below. OS, outer sulcus; PD, prosensory domain; KO, Kölliker’s organ.

(A) RNA in situ hybridization for *Etv4* and *Etv5.* The two brackets indicate *Etv4/5* expression in the outer sulcus (OS, left) and prosensory domain (PD, right; lost in *Fgf20^-/-^* and *Foxg1^Cre/+^*;*Fgfr1^flox/-^* cochleae).

(B) EdU-incorporation (green). Dashed region indicates Kölliker’s organ (KO). DAPI, nuclei (blue).

**S3 Fig.**

Quantification of length-normalized number of inner hair cells (IHCs/100 µm), outer hair cells (OHCs/100 µm), and supporting cells (SCs/100 µm) overall (along the entire cochlea; top three graphs) and in the basal, middle, and apical turns (bottom three graphs) of P0 cochleae from *Fgf20^+/-^*;*ROSA^rtTA^*(Fgf20-het), *Fgf20^+/-^*;*ROSA^rtTA^*; TRE-Fgf9-IRES-eGfp (Fgf9-OA), *Fgf20^-^/-*;*ROSA^rtTA^* (Fgf20-null), and *Fgf20^-/-^*;*ROSA^rtTA^*; TRE-Fgf9-IRES-eGfp (Fgf20-rescue) mice. Dox regimens: E13.5-E15.5, E13.5, E14.5, or E15.5. P values shown are from one-way ANOVA. * indicates p < 0.05 from Tukey’s HSD (ANOVA post-hoc); n.s., not significant. Error bars, mean ± SD. Summarized in Fig 3C.

**S4 Fig.**

(A-C) Quantification of length-normalized number of (A) inner hair cells (IHCs/100 µm), (B) outer hair cells (OHCs/100 µm), and (C) supporting cells (SCs/100 µm) in the basal, middle, and apical turns of P0 cochleae from *Sox2^+/+^*, *Sox2^Ysb/+^*, *Sox2^Ysb/Ysb^*, and *Sox2^Ysb/-^* mice. P values shown are from one-way ANOVA. * indicates p < 0.05 from Tukey’s HSD (ANOVA post-hoc); n.s., not significant. Error bars, mean ± SD.

(D) Whole mount cochlea from P0 *Sox2^Ysb/-^*mice showing presence of inner and outer hair cells (phalloidin/p75NTR) and supporting cells (Prox1/Sox2, in a different cochlea) at the basal tip. Schematic shows the location of sensory epithelium at the apical turn and basal tip of *Sox2^Ysb/-^* cochleae. Scale bar, 1 mm (whole), 100 µm (basal tip).

**S5 Fig.**

(A, B) Immunofluorescence for (A) Sox2 (red) and (B) CKDN1B (green) in sections through the basal, middle, and apical turns of E14.5 *Sox2^Ysb/+^* and *Sox2^Ysb/-^* cochleae.

(C) Immunofluorescence for Ki67 (red) on serial “mid-modiolar” sections through the E14.5 and E15.5 *Sox2^Ysb/+^* and *Sox2^Ysb/-^* cochleae. Brackets indicate prosensory domain. Nine sections through the length of the cochlear duct are labeled. See whole mount cochlear duct schematic (lower left) for relative positions of the sections.

DAPI, nuclei (blue). Scale bar, 100 µm.

**S6 Fig.**

(A) Immunofluorescence for Ki67 (red) on serial “mid-modiolar” sections through the E14.5 *Fgf20^+/-^* and *Fgf20^-/-^*cochleae. Brackets indicate prosensory domain. Nine sections through the length of the cochlear duct are labeled. See whole mount cochlear duct schematic (right) for relative positions of the sections.

(B) EdU-incorporation (green) in sections through the middle turn of E14.5 *Sox2^Ysb/+^*and *Sox2^Ysb/-^* cochleae. Dashed region indicates Kölliker’s organ (KO). Bracket indicates part of Kölliker’s organ without EdU-incorporating cells in *Sox2^Ysb/-^* cochleae.

OS, outer sulcus; PD, prosensory domain; KO, Kölliker’s organ. DAPI, nuclei (blue). Scale bar, 100 µm.

**S7 Fig.**

(A) P values from two-way ANOVA analyzing the quantification in (B-D). The two factors analyzed are *Fgf20* (*Fgf20^+/+^*, *Fgf20^+/-^*, *Fgf20^-/-^*) and *Sox2* (*Sox2^+/+^*, *Sox2^Ysb/+^*, *Sox2^Ysb/Ysb^*) gene dosage. A p value < 0.05 (yellow highlight) for *Fgf20* or *Sox2* indicates that the particular factor (independent variable) has a statistically significant effect on the measurement (dependent variable). Whereas a p value < 0.05 for Interaction indicates a statistically significant interaction between the effects of the two factors on the measurement.

(B-D) Quantification of length-normalized number of (B) inner hair cells (IHCs/100 µm), (C) outer hair cells (OHCs/100 µm), and (D) supporting cells (SCs/100 µm) in the basal, middle, and apical turns of P0 cochleae from *Fgf20^+/+^*;*Sox2^+/+^*, *Fgf20^+/+^*;*Sox2^Ysb/+^*, *Fgf20^+/-^*;*Sox2^+/+^*, *Fgf20^+/-^*;*Sox2^Ysb/+^*, *Fgf20^+/+^*;*Sox2^Ysb/Ysb^*, *Fgf20^+/-^*;*Sox2^Ysb/Ysb^*, *Fgf20^-^/-*;*Sox2^+/+^*, *Fgf20^-/-^*;*Sox2^Ysb/+^*, and *Fgf20^-/-^*;*Sox2^Ysb/Ysb^* mice. Error bars, mean ± SD.

**S8 Fig.**

Results from post-hoc Tukey’s HSD analyzing the quantification results in (B-D). Letters (L, I, J, O, P, S, T; representing each measurement in S7B-S7D Figs) indicate a statistically significant decrease (p < 0.05) when comparing the row genotype against the column genotype. L, cochlear length; I, IHCs/100 µm; O, OHCs/100 µm; S, SCs/100 µm.

**S9 Fig.**

(A) Whole mount cochlea from P0 *Fgf20^+/+^*;*Sox2^+/+^*, *Fgf20^+/-^*;*Sox2^+/+^*, *Fgf20^+/+^*;*Sox2^Ysb/+^*, and *Fgf20^+/-^*;*Sox2^Ysb/+^* mice showing inner and outer hair cells (phalloidin, green) separated by inner pillar cells (p75NTR, red). Magnifications show the basal, middle, and apical turns of the cochlea. Scale bar, 100 µm (magnifications), 1 mm (whole); arrowheads indicate ectopic inner hair cells.

(B-F) Quantification of (B) cochlear duct length, (C) total inner hair cells (IHCs) and IHCs per 100 µm of the cochlear duct, (D) total outer hair cells (OHCs) and OHCs per 100 µm, and (E) IHCs/100 µm and (F) OHCs/100 µm in the basal, middle, and apical turns at P0. P values shown are from one-way ANOVA. * indicates p < 0.05 from Tukey’s HSD (ANOVA post-hoc); n.s., not significant. Error bars, mean ± SD.

**S10 Fig.**

(A, B) Immunofluorescence for (A) Sox2 (red) and (B) CDKN1B (green) in sections through the basal, middle, and apical turns of E14.5 *Fgf20^+/-^*;*Sox2^Ysb/+^*, *Fgf20^-/-^*;*Sox2^Ysb/+^*, *Fgf20^+/-^*;*Sox2^Ysb/Ysb^*, and *Fgf20^-/-^*;*Sox2^Ysb/Ysb^* cochleae.

(C) EdU-incorporation (green) in sections through the middle turn of E14.5 *Fgf20^+/-^*;*Sox2^Ysb/+^*, *Fgf20^-/-^*;*Sox2^Ysb/+^*, *Fgf20^+/-^*;*Sox2^Ysb/Ysb^*, and *Fgf20^-/-^*;*Sox2^Ysb/Ysb^*cochleae. Dashed region indicates Kölliker’s organ (KO).

(D) Immunofluorescence for Ki67 (red) on serial “mid-modiolar” sections through the E14.5 *Fgf20^+/-^*;*Sox2^Ysb/+^*, *Fgf20^-/-^*;*Sox2^Ysb/+^*, *Fgf20^+/-^*;*Sox2^Ysb/Ysb^*, and *Fgf20^-/-^*;*Sox2^Ysb/Ysb^* cochleae. Brackets indicate prosensory domain. Nine sections through the length of the cochlear duct are labeled. See whole mount cochlear duct schematic (upper right) for relative positions of the sections.

Note: unlike in Fig 7, we have switched the placement of images from *Fgf20^-/-^*;*Sox2^Ysb/+^* and *Fgf20^+/-^*;*Sox2^Ysb/Ysb^*cochleae to facilitate comparison. OS, outer sulcus; PD, prosensory domain; KO, Kölliker’s organ. DAPI, nuclei (blue). Scale bar, 100 µm.

Author contributions
Conceptualization, L.M.Y., S.H., and D.M.O.; Methodology, L.M.Y., S.H., and D.M.O.; Formal Analysis, L.M.Y. and S.H.; Investigation: L.M.Y. and S.H.; Resources: K.S.E.C., S.H., and D.M.O.; Writing – Original Draft: L.M.Y. and D.M.O.; Writing – Review & Editing: L.M.Y., K.S.E.C., S.H., and D.M.O.; Supervision: D.M.O.; Funding Acquisition: S.H. and D.M.O.

## References

1. Basch ML, Brown RM, Jen H-I, Groves AK. Where hearing starts: the development of the mammalian cochlea. J Anat. 2016 Feb;228(2):233–54.

2. Fekete DM, Muthukumar S, Karagogeos D. Hair Cells and Supporting Cells Share a Common Progenitor in the Avian Inner Ear. J Neurosci. 1998 Oct 1;18(19):7811–21.

3. Chen P, Segil N. p27(Kip1) links cell proliferation to morphogenesis in the developing organ of Corti. Development. 1999 Apr;126(8):1581–90.

4. Lee Y-S, Liu F, Segil N. A morphogenetic wave of p27Kip1 transcription directs cell cycle exit during organ of Corti development. Development. 2006 Aug 1;133(15):2817–26.

5. Bok J, Zenczak C, Hwang CH, Wu DK. Auditory ganglion source of Sonic hedgehog regulates timing of cell cycle exit and differentiation of mammalian cochlear hair cells. Proc Natl Acad Sci U S A. 2013 Aug 20;110(34):13869–74.

6. Jahan I, Pan N, Kersigo J, Fritzsch B. Neurod1 Suppresses Hair Cell Differentiation in Ear Ganglia and Regulates Hair Cell Subtype Development in the Cochlea. Riley B, editor. PLOS ONE. 2010 Jul 22;5(7):e11661.

7. Matei V, Pauley S, Kaing S, Rowitch D, Beisel K w., Morris K, et al. Smaller inner ear sensory epithelia in Neurog1 null mice are related to earlier hair cell cycle exit. Dev Dyn. 2005 Nov 1;234(3):633–50.

8. Tateya T, Imayoshi I, Tateya I, Hamaguchi K, Torii H, Ito J, et al. Hedgehog signaling regulates prosensory cell properties during the basal-to-apical wave of hair cell differentiation in the mammalian cochlea. Development. 2013 Sep 15;140(18):3848–57.

9. Gu R, Brown RM, Hsu C-W, Cai T, Crowder AL, Piazza VG, et al. Lineage tracing of Sox2-expressing progenitor cells in the mouse inner ear reveals a broad contribution to non-sensory tissues and insights into the origin of the organ of Corti. Dev Biol. 2016 01;414(1):72–84.

10. Mak ACY, Szeto IYY, Fritzsch B, Cheah KSE. Differential and overlapping expression pattern of SOX2 and SOX9 in inner ear development. Gene Expr Patterns. 2009 Sep 1;9(6):444–53.

11. Dong S, Leung KKH, Pelling AL, Lee PYT, Tang ASP, Heng HHQ, et al. Circling, Deafness, and Yellow Coat Displayed by Yellow Submarine (Ysb) and Light Coat and Circling (Lcc) Mice with Mutations on Chromosome 3. Genomics. 2002 Jun;79(6):777–84.

12. Kiernan AE, Pelling AL, Leung KKH, Tang ASP, Bell DM, Tease C, et al. Sox2 is required for sensory organ development in the mammalian inner ear. Nature. 2005 Apr 21;434(7036):1031–5.

13. Dabdoub A, Puligilla C, Jones JM, Fritzsch B, Cheah KSE, Pevny LH, et al. Sox2 signaling in prosensory domain specification and subsequent hair cell differentiation in the developing cochlea. Proc Natl Acad Sci U S A. 2008 Nov 25;105(47):18396–401.

14. Pan W, Jin Y, Chen J, Rottier RJ, Steel KP, Kiernan AE. Ectopic Expression of Activated Notch or SOX2 Reveals Similar and Unique Roles in the Development of the Sensory Cell Progenitors in the Mammalian Inner Ear. J Neurosci. 2013 Oct 9;33(41):16146–57.

15. Puligilla C, Kelley MW. Dual role for Sox2 in specification of sensory competence and regulation of Atoh1 function. Dev Neurobiol. 2016;77(1):3–13.

16. Ebeid M, Huh S-H. FGF signaling: diverse roles during cochlear development. BMB Rep. 2017 Oct 31;50(10):487–95.

17. Hayashi T, Ray CA, Bermingham-McDonogh O. Fgf20 Is Required for Sensory Epithelial Specification in the Developing Cochlea. J Neurosci. 2008 Jun 4;28(23):5991–9.

18. Huh S-H, Warchol ME, Ornitz DM. Cochlear progenitor number is controlled through mesenchymal FGF receptor signaling. eLife. 2015 Apr 27;4:e05921.

19. Ono K, Kita T, Sato S, O’Neill P, Mak S-S, Paschaki M, et al. FGFR1-Frs2/3 Signalling Maintains Sensory Progenitors during Inner Ear Hair Cell Formation. Cheah KSE, editor. PLOS Genet. 2014 Jan 23;10(1):e1004118.

20. Pirvola U, Ylikoski J, Trokovic R, Hébert JM, McConnell SK, Partanen J. FGFR1 is required for the development of the auditory sensory epithelium. Neuron. 2002 Aug 15;35(4):671–80.

21. Huh S-H, Jones J, Warchol ME, Ornitz DM. Differentiation of the Lateral Compartment of the Cochlea Requires a Temporally Restricted FGF20 Signal. Groves A, editor. PLOS Biol. 2012 Jan 3;10(1):e1001231.

22. Hébert JM, McConnell SK. Targeting of cre to the Foxg1 (BF-1) Locus Mediates loxP Recombination in the Telencephalon and Other Developing Head Structures. Dev Biol. 2000 Jun 15;222(2):296–306.

23. Ornitz DM, Itoh N. The Fibroblast Growth Factor signaling pathway. Wiley Interdiscip Rev Dev Biol. 2015 May 1;4(3):215–66.

24. Zhang X, Ibrahimi OA, Olsen SK, Umemori H, Mohammadi M, Ornitz DM. Receptor Specificity of the Fibroblast Growth Factor Family: THE COMPLETE MAMMALIAN FGF FAMILY. J Biol Chem. 2006 Jun 9;281(23):15694–700.

25. Belteki G, Haigh J, Kabacs N, Haigh K, Sison K, Costantini F, et al. Conditional and inducible transgene expression in mice through the combinatorial use of Cre-mediated recombination and tetracycline induction. Nucleic Acids Res. 2005 Jan 1;33(5):e51–e51.

26. White AC, Xu J, Yin Y, Smith C, Schmid G, Ornitz DM. FGF9 and SHH signaling coordinate lung growth and development through regulation of distinct mesenchymal domains. Development. 2006 Apr 15;133(8):1507–17.

27. Scholzen T, Gerdes J. The Ki-67 protein: From the known and the unknown. J Cell Physiol. 2000 Mar 1;182(3):311–22.

28. Kelley MW. Cellular commitment and differentiation in the organ of Corti. Int J Dev Biol. 2007;51(6–7):571–83.

29. Benito-Gonzalez A, Doetzlhofer A. Hey1 and Hey2 Control the Spatial and Temporal Pattern of Mammalian Auditory Hair Cell Differentiation Downstream of Hedgehog Signaling. J Neurosci. 2014 Sep 17;34(38):12865–76.

30. Bermingham NA, Hassan BA, Price SD, Vollrath MA, Ben-Arie N, Eatock RA, et al. Math1: an essential gene for the generation of inner ear hair cells. Science. 1999 Jun 11;284(5421):1837–41.

31. Cai T, Seymour ML, Zhang H, Pereira FA, Groves AK. Conditional Deletion of Atoh1 Reveals Distinct Critical Periods for Survival and Function of Hair Cells in the Organ of Corti. J Neurosci. 2013 Jun 12;33(24):10110–22.

32. Woods C, Montcouquiol M, Kelley MW. Math1 regulates development of the sensory epithelium in the mammalian cochlea. Nat Neurosci. 2004 Dec;7(12):1310–8.

33. Munnamalai V, Hayashi T, Bermingham-McDonogh O. Notch Prosensory Effects in the Mammalian Cochlea Are Partially Mediated by Fgf20. J Neurosci. 2012 Sep 12;32(37):12876–84.

34. Pauley S, Lai E, Fritzsch B. Foxg1 is required for morphogenesis and histogenesis of the mammalian inner ear. Dev Dyn. 2006 Sep;235(9):2470–82.

35. Brown AS, Epstein DJ. Otic ablation of smoothened reveals direct and indirect requirements for Hedgehog signaling in inner ear development. Development. 2011 Sep 15;138(18):3967–76.

36. Kanzaki S, Beyer LA, Swiderski DL, Izumikawa M, Stöver T, Kawamoto K, et al. p27Kip1 deficiency causes organ of Corti pathology and hearing loss. Hear Res. 2006 Apr 1;214(1):28–36.

37. Ahmed M, Wong EYM, Sun J, Xu J, Wang F, Xu P-X. Eya1-Six1 Interaction Is Sufficient to Induce Hair Cell Fate in the Cochlea by Activating Atoh1 Expression in Cooperation with Sox2. Dev Cell. 2012 Feb;22(2):377–90.

38. Kempfle JS, Turban JL, Edge ASB. Sox2 in the differentiation of cochlear progenitor cells. Sci Rep. 2016;6:23293.

39. Bosman EA, Quint E, Fuchs H, Hrabé de Angelis M, Steel KP. Catweasel mice: a novel role for Six1 in sensory patch development and a model for branchio-oto-renal syndrome. Dev Biol. 2009 Apr 15;328(2):285–96.

40. Brooker R. Notch ligands with contrasting functions: Jagged1 and Delta1 in the mouse inner ear. Development. 2006 Apr 1;133(7):1277–86.

41. Kiernan AE, Ahituv N, Fuchs H, Balling R, Avraham KB, Steel KP, et al. The Notch ligand Jagged1 is required for inner ear sensory development. Proc Natl Acad Sci U S A. 2001;98(7):3873–3878.

42. Kiernan AE, Xu J, Gridley T. The Notch ligand JAG1 is required for sensory progenitor development in the mammalian inner ear. PLOS Genet. 2006;2(1):e4.

43. Kiernan AE, Li R, Hawes NL, Churchill GA, Gridley T. Genetic background modifies inner ear and eye phenotypes of jag1 heterozygous mice. Genetics. 2007 Sep;177(1):307–11.

44. Moayedi Y, Basch ML, Pacheco NL, Gao SS, Wang R, Harrison W, et al. The Candidate Splicing Factor Sfswap Regulates Growth and Patterning of Inner Ear Sensory Organs. Wu DK, editor. PLOS Genet. 2014 Jan 2;10(1):e1004055.

45. Tsai H, Hardisty RE, Rhodes C, Kiernan AE, Roby P, Tymowska-Lalanne Z, et al. The mouse slalom mutant demonstrates a role for Jagged1 in neuroepithelial patterning in the organ of Corti. Hum Mol Genet. 2001 Mar 1;10(5):507–12.

46. Yamamoto N, Chang W, Kelley MW. Rbpj regulates development of prosensory cells in the mammalian inner ear. Dev Biol. 2011 May;353(2):367–79.

47. Basch ML, Ohyama T, Segil N, Groves AK. Canonical Notch Signaling Is Not Necessary for Prosensory Induction in the Mouse Cochlea: Insights from a Conditional Mutant of RBPj. J Neurosci. 2011 Jun 1;31(22):8046–58.

48. Kopecky BJ, Jahan I, Fritzsch B. Correct Timing of Proliferation and Differentiation is Necessary for Normal Inner Ear Development and Auditory Hair Cell Viability: Mycs’ Role in Ear Development. Dev Dyn. 2013 Feb;242(2):132–47.

49. Chen P, Johnson JE, Zoghbi HY, Segil N. The role of Math1 in inner ear development: Uncoupling the establishment of the sensory primordium from hair cell fate determination. Development. 2002 May;129(10):2495–505.

50. Haque K, Pandey AK, Zheng H-W, Riazuddin S, Sha S-H, Puligilla C. MEKK4 Signaling Regulates Sensory Cell Development and Function in the Mouse Inner Ear. J Neurosci. 2016 Jan 27;36(4):1347–61.

51. Trokovic N, Trokovic R, Mai P, Partanen J. Fgfr1 regulates patterning of the pharyngeal region. Genes Dev. 2003 Jan 1;17(1):141–53.

52. Hayashi S, Lewis P, Pevny L, McMahon AP. Efficient gene modulation in mouse epiblast using a Sox2Cre transgenic mouse strain. Mech Dev. 2002 Dec;119 Suppl 1:S97–101.

53. Shaham O, Smith AN, Robinson ML, Taketo MM, Lang RA, Ashery-Padan R. Pax6 is essential for lens fiber cell differentiation. Development. 2009 Aug 1;136(15):2567–78.

54. Avinash GB, Nuttall AL, Raphael Y. 3-D analysis of F-actin in stereocilia of cochlear hair cells after loud noise exposure. Hear Res. 1993 May 1;67(1):139–46.

55. Mueller KL, Jacques BE, Kelley MW. Fibroblast Growth Factor Signaling Regulates Pillar Cell Development in the Organ of Corti. J Neurosci. 2002 Nov 1;22(21):9368–77.

56. Bermingham-McDonogh O, Oesterle EC, Stone JS, Hume CR, Huynh HM, Hayashi T. Expression of Prox1 during mouse cochlear development. J Comp Neurol. 2006 May 10;496(2):172–86.

