## Supplemental data for "Sox2 and FGF20 interact to regulate organ of Corti hair cell and supporting cell development in a spatially-graded manner"

S1 Fig

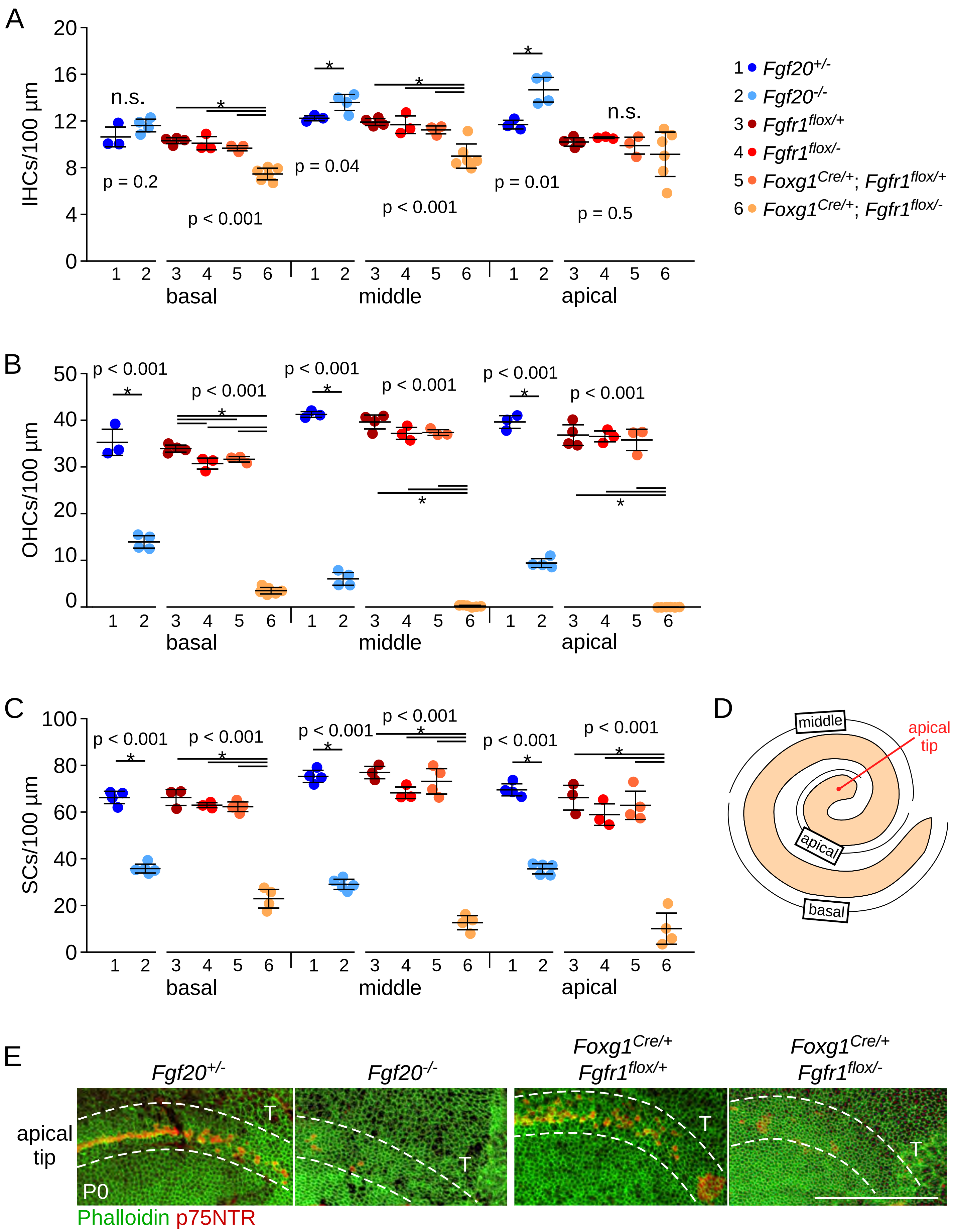

A
IHCs/100 µm
n.s.
p = 0.2
p < 0.001
p = 0.04
p < 0.001
p = 0.01
n.s.
p = 0.5
basal
middle
apical
1 Fgf20+/-
2 Fgf20-/-
3 Fgf1flox/+
4 Fgfr1flox/-
5 Foxg1Cre/+; Fgfr1flox/+
6 Foxg1Cre/+; Fgfr1flox/-
B
OHCs/100 µm
C
SCs/100 µm
D
middle
apical tip
apical
basal
E
Fgf20+/-
Fgf20-/-
Foxg1Cre/+ Fgfr1flox/+
Foxg1Cre/+ Fgfr1flox/-
apical tip
P0
Phalloidin p75NTR

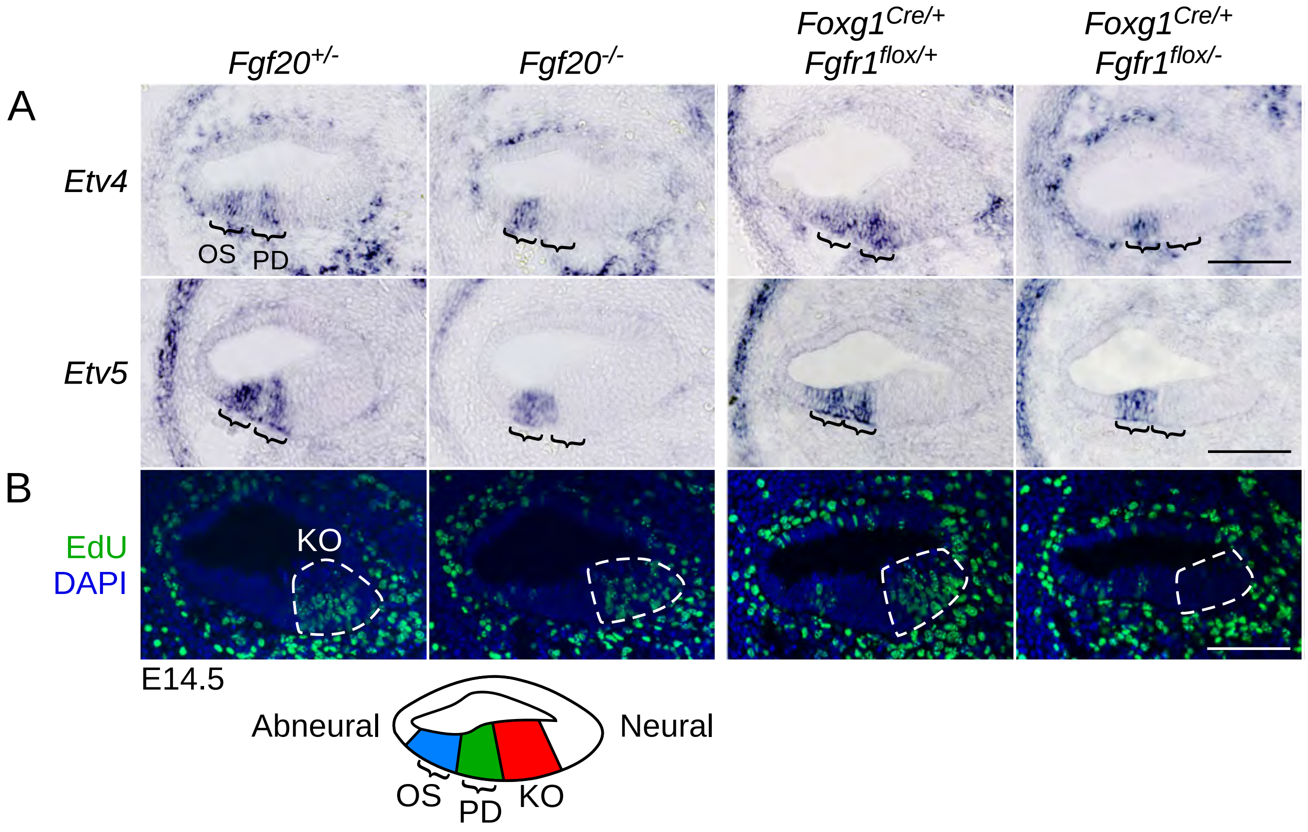
Fgf20+/-
Fgf20-/-
Foxg1Cre/+ Fgfr1flox/+
Foxg1Cre/+ Fgfr1flox/-
A
Etv4
OS
PD
Etv5
B
EdU
DAPI
KO
E14.5
Abneural
Neural
OS
PD
KO

S3 Fig

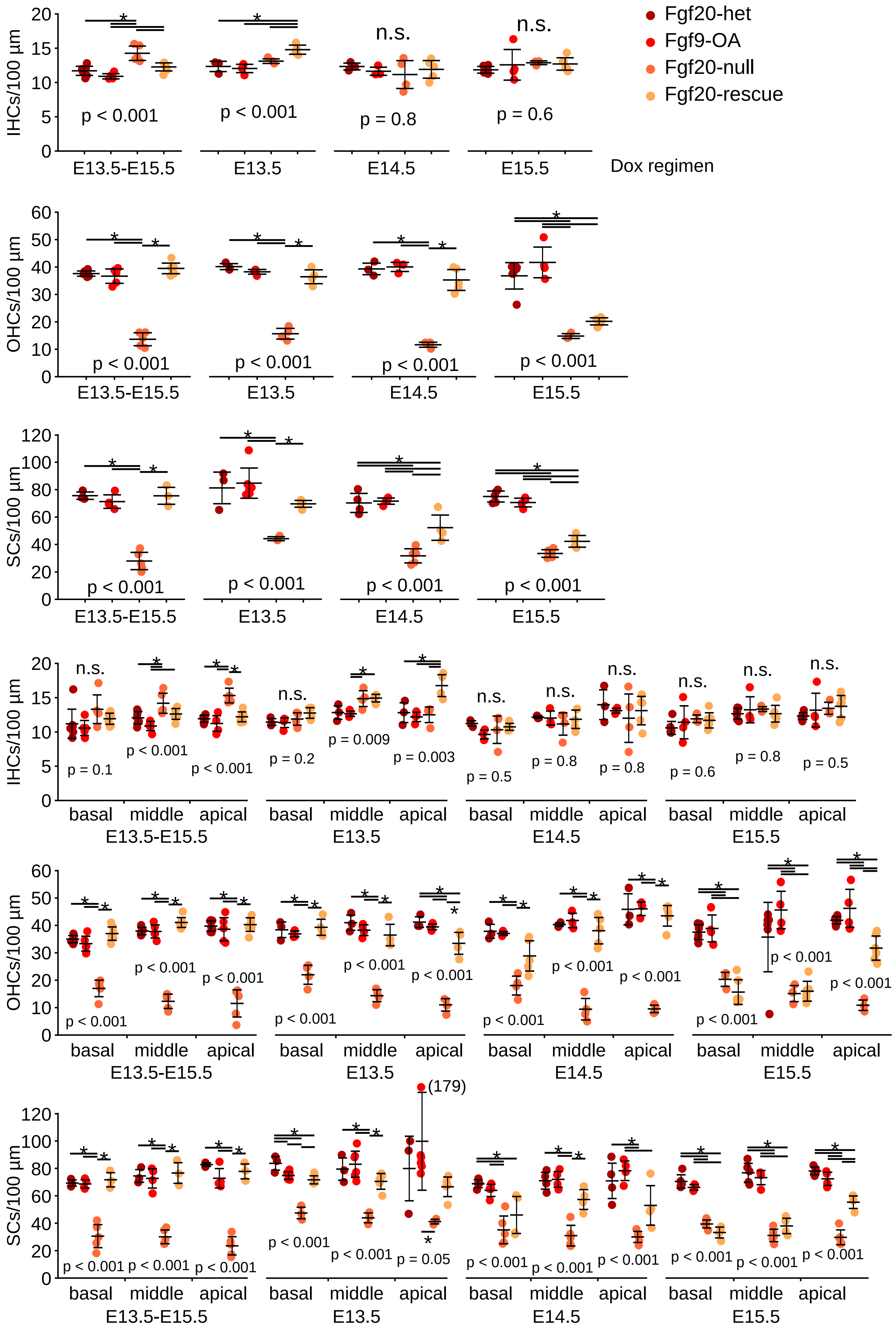

Fgf20-het
Fgf9-OA
Fgf20-null
Fgf20-rescue
IHCs/100 µm
n.s.
n.s.
p < 0.001
p < 0.001
p = 0.8
p = 0.6
E13.5-E15.5
E13.5
E14.5
E15.5
Dox regimen
OHCs/100 µm
p < 0.001
p < 0.001
p < 0.001
p < 0.001
SCs/100 µm
p < 0.001
p < 0.001
p < 0.001
p < 0.001
IHCs/100 µm
p = 0.1
p < 0.001
p < 0.001
p = 0.2
p = 0.009
p = 0.003
p = 0.5
p = 0.8
p = 0.8
p = 0.6
p = 0.8
p = 0.5
basal
middle
apical
OHCs/100 µm
SCs/100 µm
(179)
p = 0.05

S4 Fig

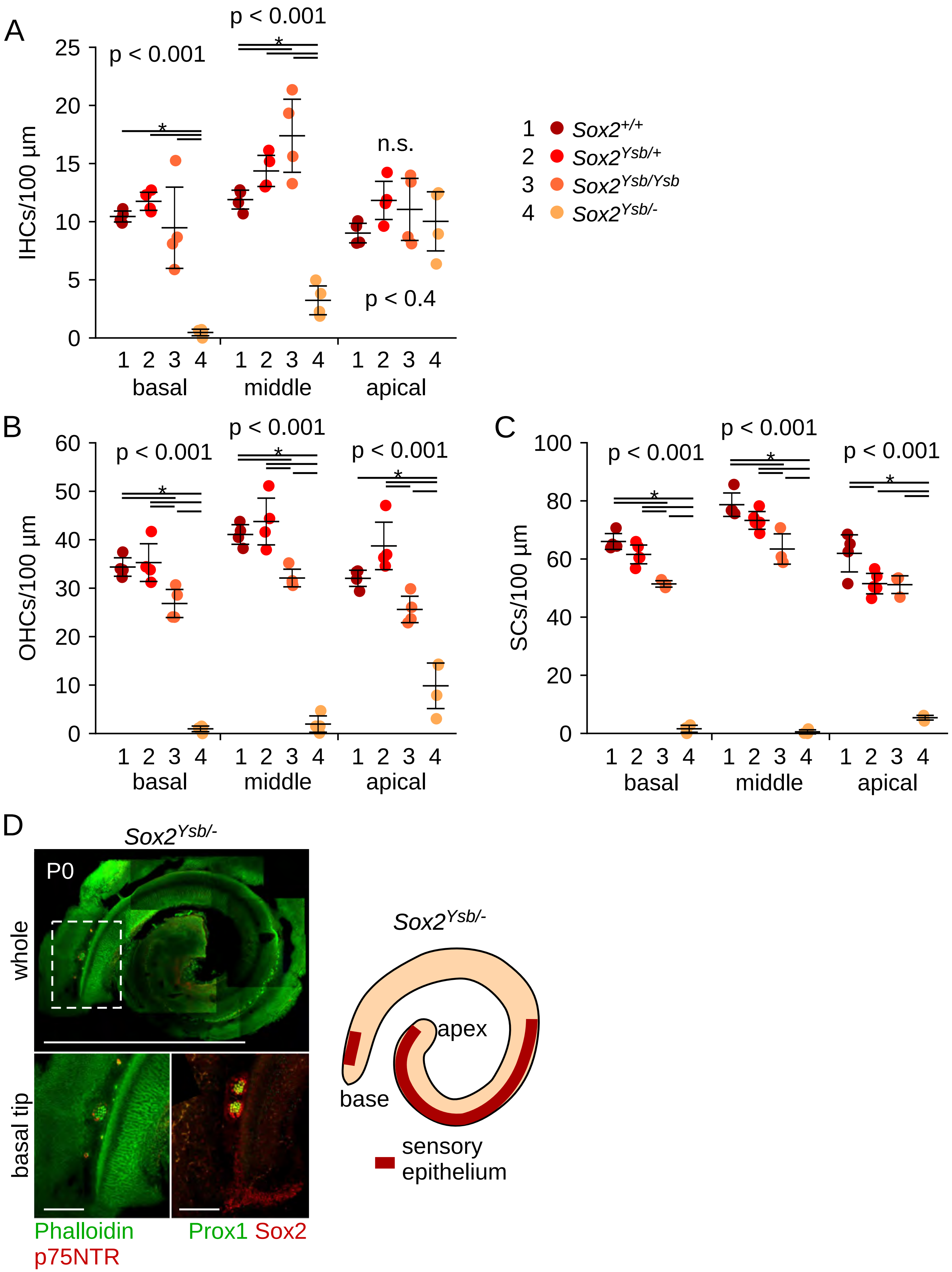

A
IHCs/100 µm
p < 0.001
p < 0.001
n.s.
p < 0.4
1 2 3 4
basal
middle
apical
1 Sox2+/+
2 Sox2Ysb/+
3 Sox2Ysb/Ysb
4 Sox2Ysb/-
B
OHCs/100 µm
C
SCs/100 µm
D
Sox2Ysb/-
P0
whole
basal tip
Phalloidin
p75NTR
Prox1 Sox2
apex
base
sensory
epithelium

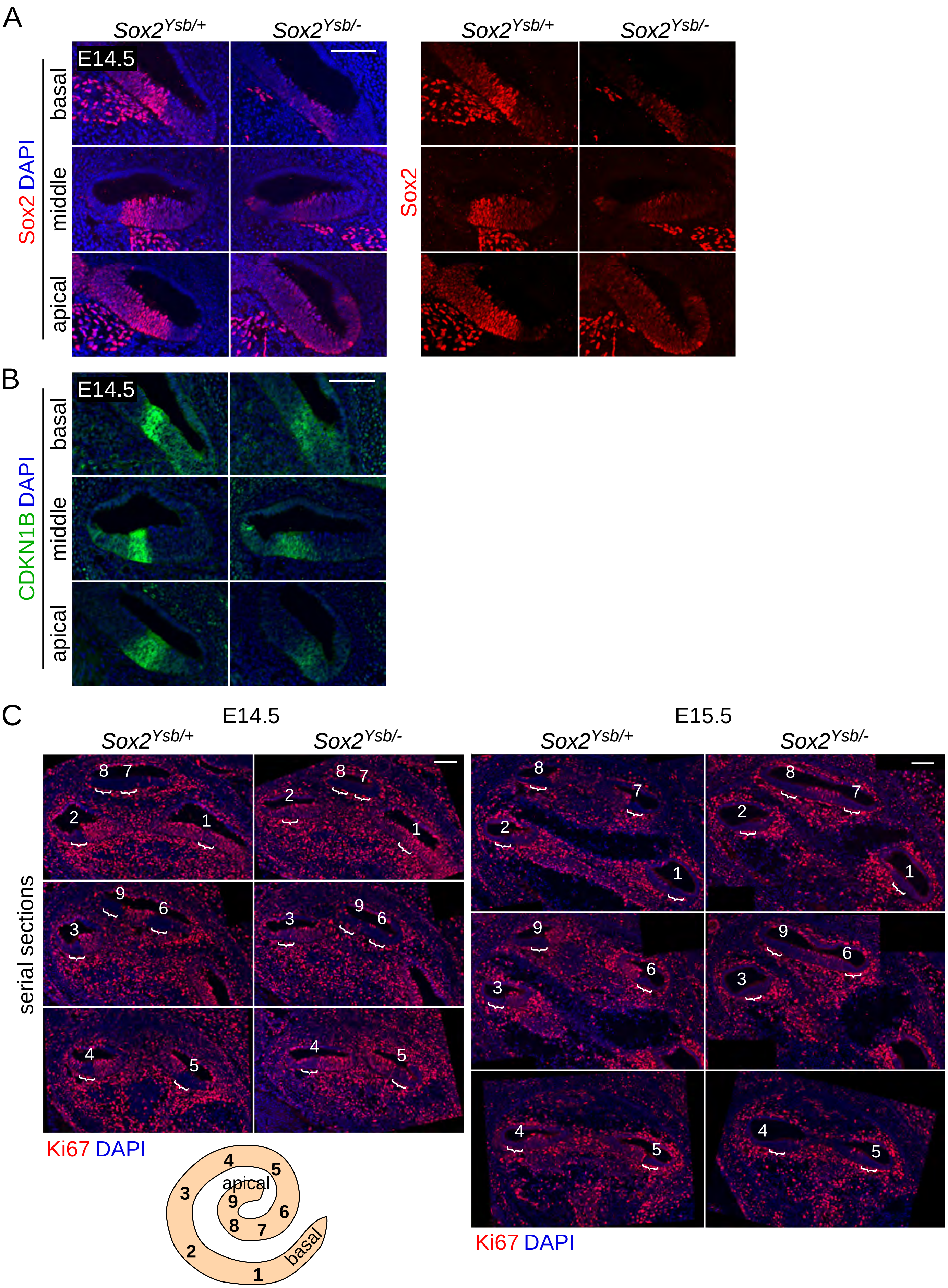
A
Sox2Ysb/+
Sox2Ysb/-
E14.5
Sox2 DAPI
basal
middle
apical
Sox2
B
E14.5
CDKN1B DAPI
basal
middle
apical
C
E14.5
E15.5
Sox2Ysb/+
Sox2Ysb/-
serial sections
Ki67 DAPI
apical
basal
1
2
3
4
5
6
7
8
9

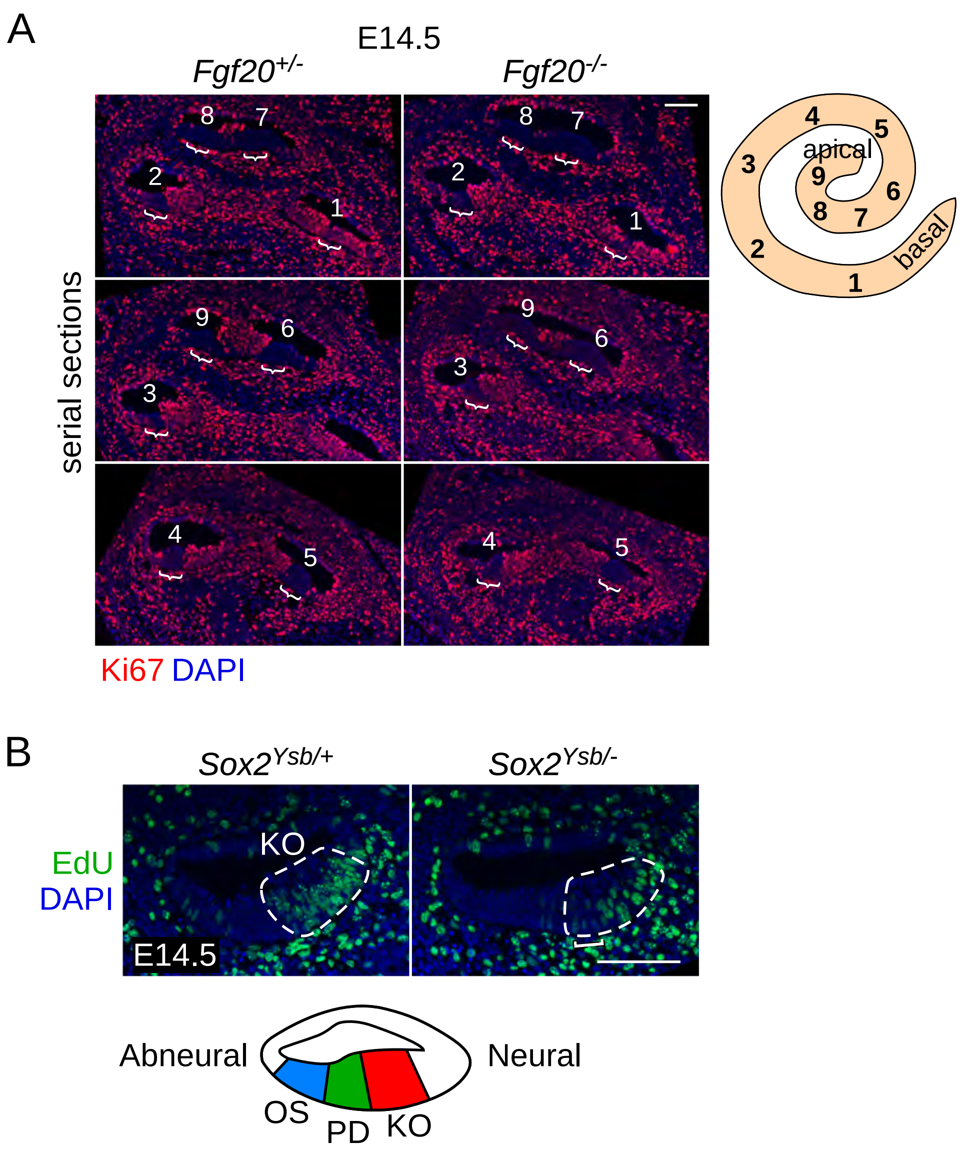
A
E14.5
Fgf20+/-
Fgf20-/-
serial sections
8
7
2
1
9
6
3
4
5
apical
basal
Ki67 DAPI
B
Sox2Ysb/+
Sox2Ysb/-
EdU
DAPI
KO
E14.5
Abneural
Neural
OS
PD
KO

A

| 2-way ANOVA | *Fgf20* | | | *Sox2* | | | Interaction | | |
|---|---|---|---|---|---|---|---|---|---|
| | basal | middle | apical | basal | middle | apical | basal | middle | apical |
| IHC | $p < 0.001$ | $p < 0.001$ | $p = 0.2$ | $p < 0.001$ | $p = 0.2$ | $p = 0.01$ | $p < 0.001$ | $p < 0.001$ | $p < 0.001$ |
| OHC | $p < 0.001$ | $p < 0.001$ | $p < 0.001$ | $p < 0.001$ | $p < 0.001$ | $p < 0.001$ | $p = 0.002$ | $p = 0.002$ | $p < 0.001$ |
| SC | $p < 0.001$ | $p < 0.001$ | $p < 0.001$ | $p < 0.001$ | $p < 0.001$ | $p < 0.001$ | $p = 0.01$ | $p = 0.01$ | $p = 0.7$ |

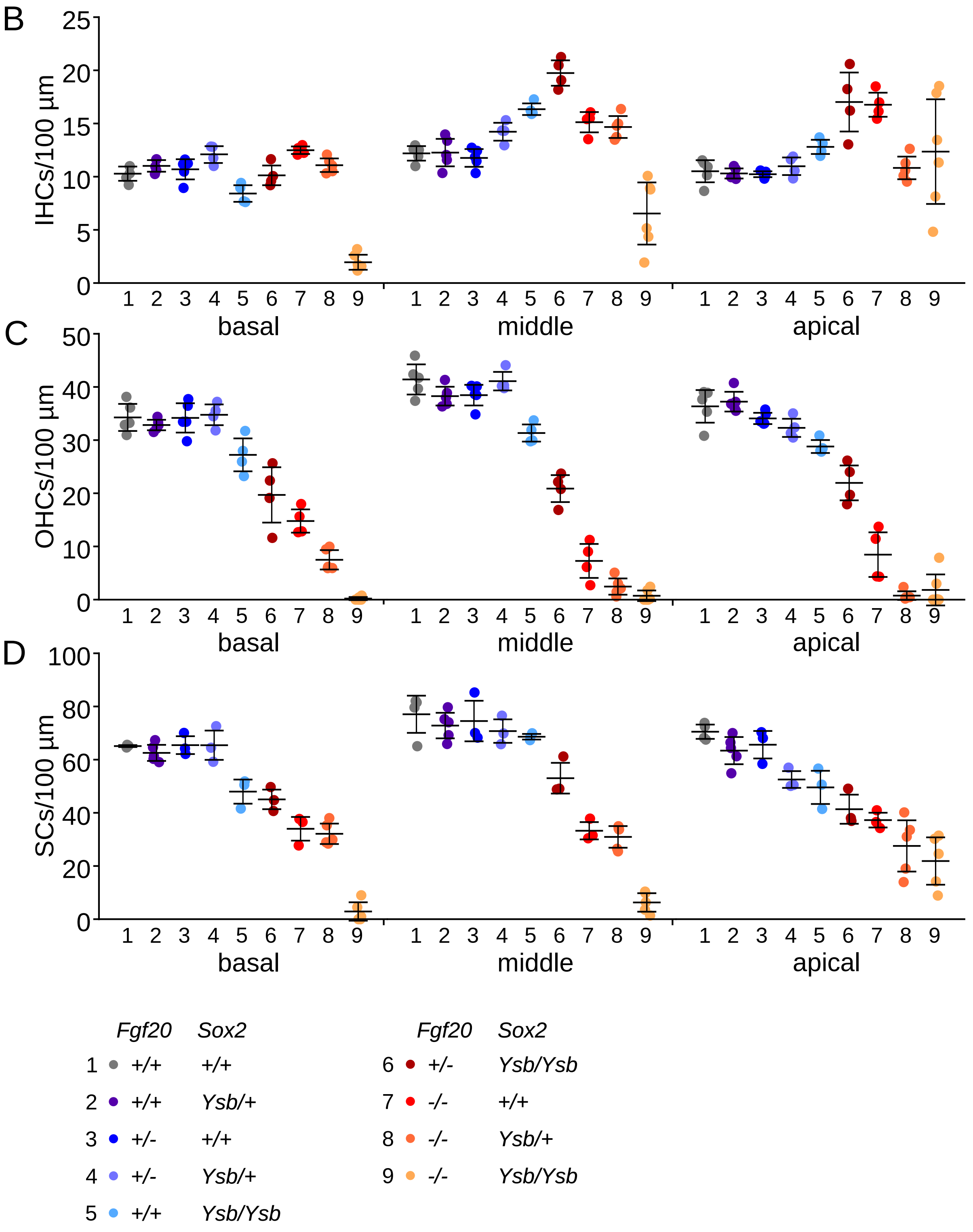

S8 Fig

| basal | | | 1<br>+/+<br>+/+ | 2<br>+/+<br>Ysb/+ | 3<br>+/-<br>+/+ | 4<br>+/-<br>Ysb/+ | 5<br>+/+<br>Ysb/Ysb | 6<br>+/-<br>Ysb/Ysb | 7<br>-/-<br>+/+ | 8<br>-/-<br>Ysb/+ | 9<br>-/-<br>Ysb/Ysb |
|---|---|---|---|---|---|---|---|---|---|---|---|
| | *Fgf20* | *Sox2* | | | | | | | | | |
| 1 | +/+ | +/+ | | | | I | | | I | | |
| 2 | +/+ | Ysb/+ | | | | | | | | | |
| 3 | +/- | +/+ | | | | | | | | | |
| 4 | +/- | Ysb/+ | | | | | | | | | |
| 5 | +/+ | Ysb/Ysb | I O S | I S | I O S | I O S | | | I | I | |
| 6 | +/- | Ysb/Ysb | O S | O S | O S | I O S | O | | I | | |
| 7 | -/- | +/+ | O S | O S | O S | O S | O S | | | | |
| 8 | -/- | Ysb/+ | O S | O S | O S | O S | O S | O S | O | | |
| 9 | -/- | Ysb/Ysb | I O S | I O S | I O S | I O S | I O S | I O S | I O S | I O S | |

| middle | | | 1<br>+/+<br>+/+ | 2<br>+/+<br>Ysb/+ | 3<br>+/-<br>+/+ | 4<br>+/-<br>Ysb/+ | 5<br>+/+<br>Ysb/Ysb | 6<br>+/-<br>Ysb/Ysb | 7<br>-/-<br>+/+ | 8<br>-/-<br>Ysb/+ | 9<br>-/-<br>Ysb/Ysb |
|---|---|---|---|---|---|---|---|---|---|---|---|
| | *Fgf20* | *Sox2* | | | | | | | | | |
| 1 | +/+ | +/+ | | | | | I | I | | | |
| 2 | +/+ | Ysb/+ | | | | | I | I | | | |
| 3 | +/- | +/+ | | | | | I | I | | | |
| 4 | +/- | Ysb/+ | | | | | | I | | | |
| 5 | +/+ | Ysb/Ysb | O | O | O | O | | | | | |
| 6 | +/- | Ysb/Ysb | O S | O S | O S | O S | O | | | | |
| 7 | -/- | +/+ | O S | O S | O S | O S | O S | I O S | | | |
| 8 | -/- | Ysb/+ | O S | O S | O S | O S | O S | I O S | | | |
| 9 | -/- | Ysb/Ysb | I O S | I O S | I O S | I O S | I O S | I O S | I O S | I S | |

| apical | | | 1<br>+/+<br>+/+ | 2<br>+/+<br>Ysb/+ | 3<br>+/-<br>+/+ | 4<br>+/-<br>Ysb/+ | 5<br>+/+<br>Ysb/Ysb | 6<br>+/-<br>Ysb/Ysb | 7<br>-/-<br>+/+ | 8<br>-/-<br>Ysb/+ | 9<br>-/-<br>Ysb/Ysb |
|---|---|---|---|---|---|---|---|---|---|---|---|
| | *Fgf20* | *Sox2* | | | | | | | | | |
| 1 | +/+ | +/+ | | | | | | I | I | | |
| 2 | +/+ | Ysb/+ | | | | | | I | I | | |
| 3 | +/- | +/+ | | | | | | I | I | | |
| 4 | +/- | Ysb/+ | | | | | | I | I | | |
| 5 | +/+ | Ysb/Ysb | O S | O | | | | | | | |
| 6 | +/- | Ysb/Ysb | O S | O S | O S | O | O | | | | |
| 7 | -/- | +/+ | O S | O S | O S | O | O | O | | | |
| 8 | -/- | Ysb/+ | O S | O S | O S | O S | O S | I O | I O | | |
| 9 | -/- | Ysb/Ysb | O S | O S | O S | O S | O S | O S | O | | |

I IHCs/100 µm
O OHCs/100 µm
S SCs/100 µm

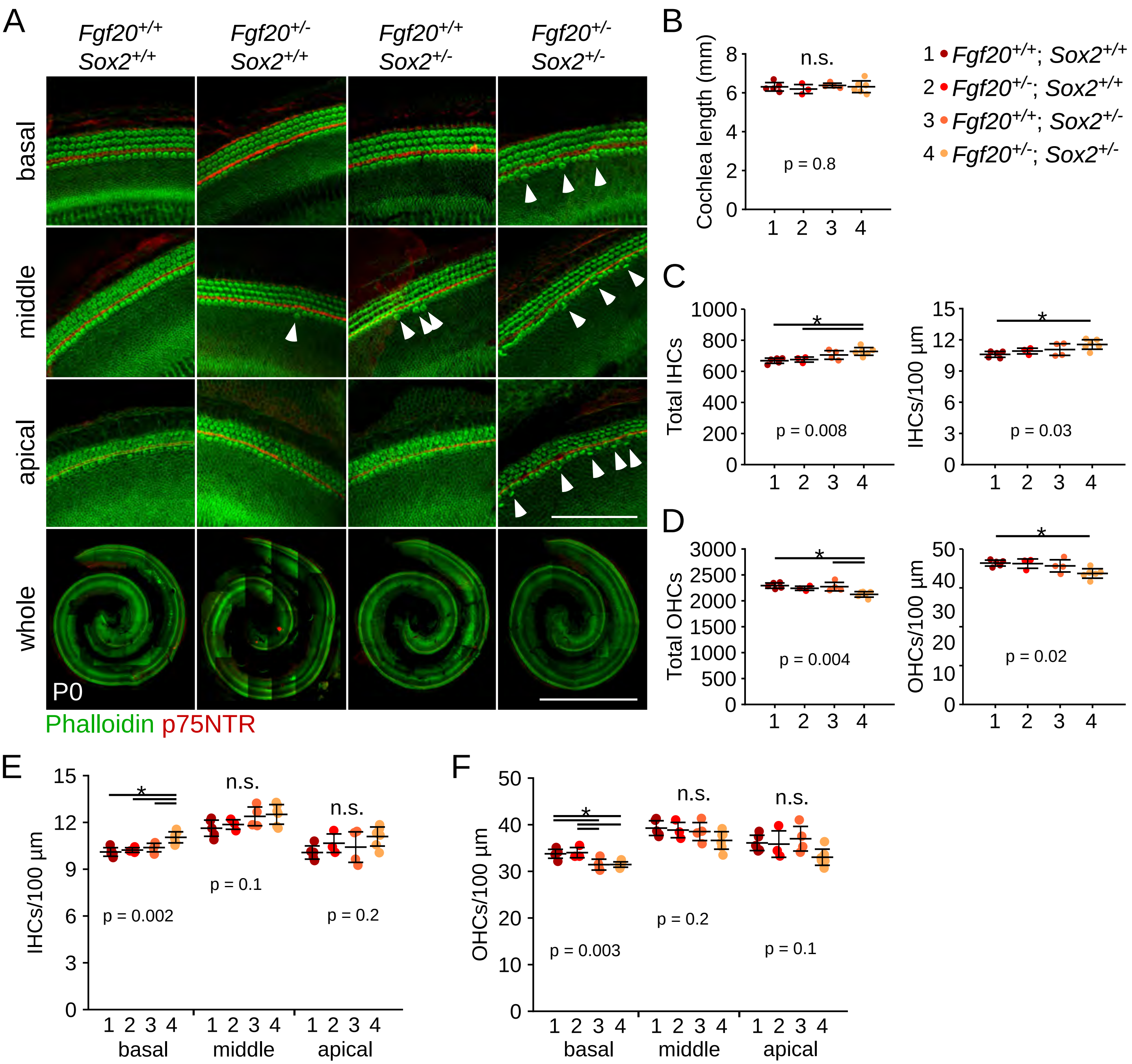

A
Fgf20+/+ Sox2+/+
Fgf20+/- Sox2+/+
Fgf20+/+ Sox2+/-
Fgf20+/- Sox2+/-
basal
middle
apical
whole
P0
Phalloidin p75NTR
B
Cochlea length (mm)
n.s.
p = 0.8
1 Fgf20+/+; Sox2+/+
2 Fgf20+/-; Sox2+/+
3 Fgf20+/+; Sox2+/-
4 Fgf20+/-; Sox2+/-
C
Total IHCs
p = 0.008
IHCs/100 µm
p = 0.03
D
Total OHCs
p = 0.004
OHCs/100 µm
p = 0.02
E
IHCs/100 µm
p = 0.002
n.s.
p = 0.1
n.s.
p = 0.2
basal
middle
apical
F
OHCs/100 µm
p = 0.003
n.s.
p = 0.2
n.s.
p = 0.1
basal
middle
apical

S10 Fig

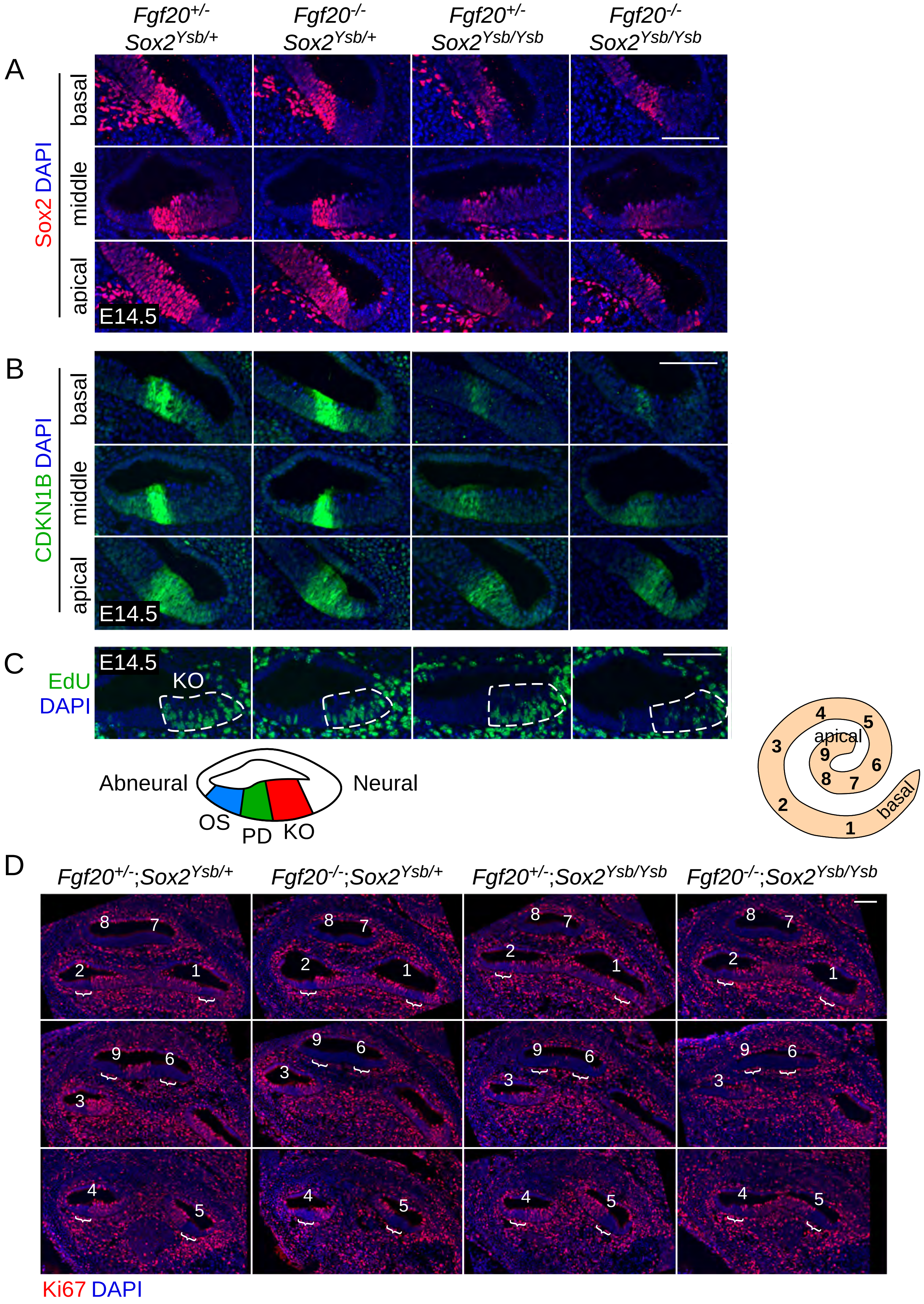

A
Fgf20+/- Sox2Ysb/+
Fgf20-/- Sox2Ysb/+
Fgf20+/- Sox2Ysb/Ysb
Fgf20-/- Sox2Ysb/Ysb
Sox2 DAPI
basal
middle
apical
E14.5
B
CDKN1B DAPI
basal
middle
apical
E14.5
C
EdU
DAPI
E14.5
KO
Abneural
Neural
OS
PD
KO
4
5
apical
3
9
6
8
7
2
basal
1
D
Fgf20+/-;Sox2Ysb/+
Fgf20-/-;Sox2Ysb/+
Fgf20+/-;Sox2Ysb/Ysb
Fgf20-/-;Sox2Ysb/Ysb
8
7
2
1
9
6
3
4
5
Ki67 DAPI
